# Zika virus noncoding RNA cooperates with the viral protein NS5 to inhibit STAT1 phosphorylation and facilitate viral pathogenesis

**DOI:** 10.1101/2021.05.18.444753

**Authors:** Andrii Slonchak, Xiaohui Wang, Harman Chaggar, Julio Aguado, Morgan Freney, Kexin Yan, Francisco J Torres, Alberto A Amarilla, Rickyle Balea, Julian D. J. Sng, Yin Xiang Setoh, Nias Peng, Daniel Watterson, Ernst Wolvetang, Andreas Suhrbier, Alexander A Khromykh

## Abstract

Zika virus (ZIKV) is a re-emerging pathogenic flavivirus, which causes microcephaly in infants and poses a continuing threat to public health. ZIKV, like all other flaviviruses, produces highly abundant noncoding RNA known as subgenomic flaviviral RNA (sfRNA). Herein we utilized wild-type and mutant ZIKV defective in production of sfRNA to elucidate for the first time how production of sfRNA affects all aspects of ZIKV pathogenesis. We found that in mouse pregnancy model of infection sfRNA is required for trans-placental dissemination of ZIKV and subsequent infection of fetal brain. Using human brain organoids, we showed that sfRNA promotes apoptosis of neural progenitor cells leading to profound cytopathicity and disintegration of organoids. We also found by transcriptome profiling and gene network analysis that in infected human placental cells sfRNA inhibits multiple antiviral pathways and promotes apoptosis with STAT1 identified as a key shared factor linking these two interconnected sfRNA activities. We further showed for the first time that sfRNA inhibits phosphorylation and nuclear translocation of STAT1 by a novel mechanism which involves binding to and stabilizing viral protein NS5. This allows accumulation of NS5 at the levels required for efficient inhibition of STAT1 phosphorylation. Thus, we elucidated the molecular mechanism by which ZIKV sfRNA exerts its functions in vertebrate hosts and discovered a co-operation between viral noncoding RNA and a viral protein as a novel strategy employed by viruses to counteract antiviral responses.

## Introduction

Flaviviruses are small enveloped viruses with single-stranded positive sense RNA genomes^1^. A unique feature of flavivirus infection is the production of noncoding RNA derived from viral 3’-untranslated regions (3’UTRs), which accumulates in infected cells in high abundance^2^. This RNA, termed subgenomic flaviviral RNA (sfRNA), is generated because of highly conserved RNA elements in the 3’UTRs that resist degradation by the cellular 5’->3’ exoribonuclease XRN-1^3, 4^. Previously, sfRNA was shown to facilitate replication and pathogenesis of West Nile (WNV)^5^ and dengue (DENV)^6–8^ viruses. However, the molecular mechanism that underpins this activity is currently not well established^9^. Furthermore, the role of sfRNA in the pathogenesis of other flaviviruses, including Zika virus (ZIKV) remains poorly defined.

Zika virus (ZIKV) is a mosquito-borne pathogenic flavivirus capable of causing re-occurring severe outbreaks in human populations^10^. Amongst flaviviruses, ZIKV is rather unique in its ability to disseminate through the placenta and establish infection in the foetal brain, resulting in neurodevelopmental abnormalities^11^. Infected mothers subsequently have a high risk of giving birth to infants with microcephaly and other neurological disorders known collectively as congenital Zika syndrome^12^. The 3’UTR of ZIKV contains two XRN-1 resistant RNA elements (xrRNAs) and is processed into two sfRNAs that differ in length^13^. To determine the role of these RNAs in ZIKV infection, we recently generated sfRNA-deficient mutant viruses and used them to show that sfRNA facilitates ZIKV transmission by mosquitoes with a mechanism involving inhibition of apoptosis in mosquito tissues leading to increased virus dissemination and secretion into saliva^14^. Herein, we used this loss-of-function system to investigate the role of ZIKV sfRNA during infection of the vertebrate host.

Using a combination of *in vivo, ex vivo* and *in cellulo* approaches, we demonstrated that sfRNA facilitates replication and pathogenesis of ZIKV and is required for virus-induced cytopathic effects. Importantly, in pregnant mice ZIKV sfRNA was crucial for establishing infection in placenta and virus dissemination into the fetal brain. Moreover, in 3D human brain organoids, ZIKV sfRNA facilitated apoptosis of infected neural progenitors, which resulted in growth retardation and eventual death of organoids. This for the first time revealed the requirement of sfRNA for neurovirulence and neuropathogenicity of ZIKV. Furthermore, in human placental cells, transcriptome profiling and pathway enrichment analysis identified the inhibitory effect of ZIKV sfRNA on multiple signaling pathways. In addition to suppressing negative regulators of apoptosis, sfRNA was found to inhibit signaling from type I, II and III interferons, as well as pro-inflammatory pathways. The subsequent gene network analysis of infected human placental cells revealed that Signal Transducer and Activator of Transcription 1 (STAT1) is a critical shared component of the identified sfRNA-affected pathways. Further experiments revealed that production of sfRNA caused a dramatic reduction in STAT1 phosphorylation and thus suppressed antiviral signalling by type I, II, and III interferons (IFN). This is the first time STAT1 is identified as a target of sfRNA. Mechanistically, we showed that Inhibition of STAT1 phosphorylation by sfRNA required cooperation between sfRNA and the viral protein NS5, whereby sfRNA bound to and stabilized NS5 leading to accumulation of NS5 in the amounts required for inhibition of STAT1 phosphorylation. Hence, we concluded that stabilization of NS5 by sfRNA is the key mechanism by which sfRNA exerts its functions in immune evasion and pathogenesis of ZIKV.

## Results

### ZIKV sfRNA facilitates virus replication, determines cytopathic effect and promotes viral pathogenesis

To elucidate the functions of ZIKV sfRNA, we previously introduced point mutations (xrRNA1’ and xrRNA2’) into each and both xrRNAs within the 3’UTR of the African ZIKV strain, ZIKV_MR766_^14^. In mosquito cells, individual mutations in xrRNA1 and xrRNA2 abrogated production of sfRNA-1 and sfRNA-2, respectively, while virus containing mutations in both xrRNAs was not viable^14^. To determine whether the phenotypes of xrRNA1’ and xrRNA2’ mutants are preserved in a vertebrate host, Vero cells were infected with wild-type (WT) ZIKV and the two viral mutants. The interferon-deficient Vero cells^15^ were used in this experiment because they support replication of all three viruses at comparable levels^14^. Unlike in insect cells^14^, the mutation in xrRNA2 not only abolished production of sfRNA-2, but also dramatically reduced accumulation of sfRNA-1 in mammalian cells (Fig 1A), although they contained similar levels of viral genomic RNA (Fig 1B). As the mutation in xrRNA2 largely impairs production of both sfRNAs in vertebrate cells without compromising virus viability, the xrRNA2’ ZIKV mutant can be used to assess the effect of nearly complete sfRNA deficiency on virus replication and pathogenesis. In turn, the xrRNA1’ mutant, which is deficient in sfRNA-1 only, can be used to study the impact of partial sfRNA-deficiency on the virus.

**Figure 1.**
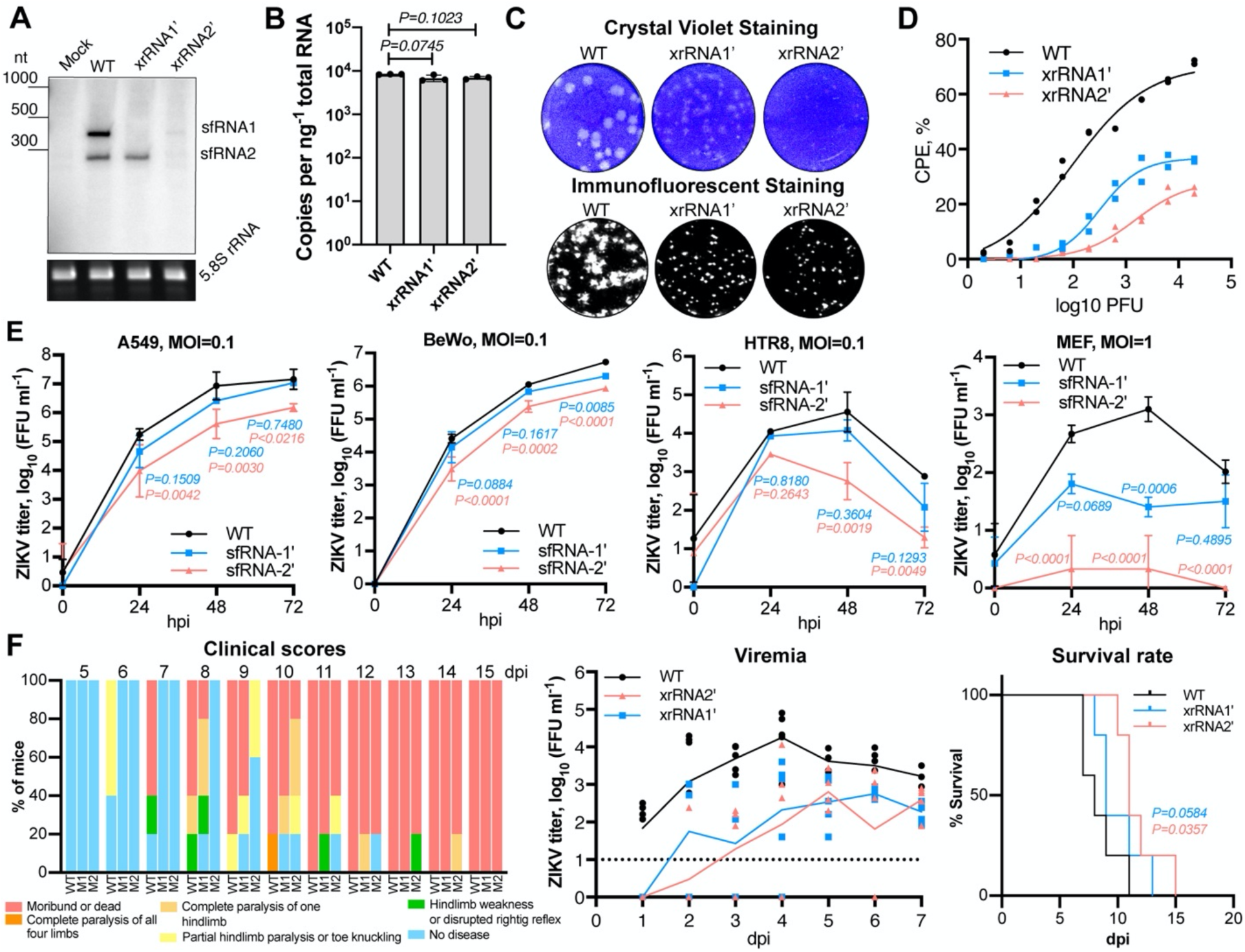
ZIKV sfRNA facilitates viral replication, cytotoxicity and pathogenesis in mammalian host. **(A)** Detection of ZIKV sfRNA in Vero cells infected with WT, xrRNA1’ and xrRNA2’ viruses. Cells were infected at MOI=1, RNA was isolated at 72hpi and used for Northern blotting. Blot was hybridised with a probe complementary to the most terminal 31nt of viral genome. Bottom panel shows rRNA visualised by Et-Br staining as a loading control. The blot is a representative image from three independent experiments that showed similar results. **(B)** Quantification of viral RNA in samples used for northern blotting in (A). RNA copy numbers were determined by qRT-PCR with normalisation to input RNA amount. The values are the means from three biological replicates ±SD, statistical analysis is by one-way ANOVA. **(C)** ZIKV plaque morphology on a monolayer of Vero cells at 72hpi. Bottom panel demonstrates virus replication foci visualised by immunostaining of Vero cells inoculated with the same virus samples. **(D)** Cytotoxicity of WT and sfRNA-deficient ZIKV mutants determined using Viral ToxGlo Assay. Vero cells were infected at the indicated viral dose per well and CPE was measured at 72hpi. %CPE is calculated with the reference to uninfected cells grown for the same period of time. Individual values from two independent experiments with technical triplicates in each are displayed. **(E)** ZIKV titres in culture fluids of human and mouse cells infected with WT and sfRNA deficient ZIKV. Cells were infected at MOI=0.1. Values are the means from three independent experiments ±SD. Statistical analysis is by two-way ANOVA with Dunnett correction. **(F)** Replication of WT and sfRNA- deficient ZIKV in AG129 mice. Mice were inoculated with 10^4^ FFU of the viruses by subcutaneous injection and were monitored for disease symptoms twice a day for 15 days and blood was collected daily via tail bleeding. Mean values were used to build the curves in viremia analysis and all individual values are shown on the plot. The results of the statistical analysis for viremia are summarised in Supplementary table 1, survival rates were compared using Gehan-Breslow-Wilcoxon test. Viral titres were determined by foci-forming immunoassay with cross-reactive mouse anti-E protein monoclonal antibodies (4G2) as presented in (E) and human anti-E protein (Z67) as presented in (F).

To elucidate the functions of ZIKV sfRNAs, the effect of sfRNA deficiency on viral cytotoxicity and replication in cultured cells was first assessed. Plaque-assay on Vero cells demonstrated that WT ZIKV was capable of forming large clear plaques, whereas plaques formed by xrRNA1’ mutant had reduced size and were less clear. No plaques were clearly visible in cells infected with xrRNA2’ mutant (Fig 1C, top row), while the foci-forming immunoassay showed clearly visible virus-induced foci (Fig 1C, bottom row) indicative of established ZIKV infection. The quantitative viral cytopathic effect (CPE) assay revealed up to 3-fold higher levels of CPE in Vero cells infected with WT virus, when compared with the xrRNA2’ mutant (Fig 1D). Together these results demonstrated that sfRNAs are required for the CPE induced by ZIKV infection in vertebrate cells. Given that WT and mutant viruses replicate in Vero cells at the same level (Fig 1B), the reduced cytopathic properties of sfRNA-deficient ZIKV cannot be attributed to the lower viral load. Therefore, the sfRNA likely has a specific function in promoting cell death.

In IFN response-competent human cell lines A549, BeWo, and HTR8 infected at MOI=0.1, the xrRNA1’ mutant ZIKV replicated with efficiency comparable to WT virus, whereas the replication of xrRNA2’ mutant was highly attenuated (Fig 1E). In IFN response-competent mouse embryonic fibroblasts (MEF), both mutants were incapable of productive infection after virus inoculation at MOI=0.1 (Supplementary Figure 1). At MOI=1, the xrRNA1’ mutant established infection, although at a lower level than WT virus, whereas replication of xrRNA2’ was still barely detectable (Fig 1E). These results demonstrated that production of sfRNAs is required for efficient replication of ZIKV in IFN response-competent mammalian cells.

To determine if sfRNAs contribute to pathogenesis of ZIKV infection *in vivo,* AG129 mice were used as an infection model. These mice are deficient in receptors for IFN-*α*/*β* and IFN-*γ*, and, unlike WT mice, support replication of ZIKV and develop disease^16^. Mice infected with sfRNA-deficient mutants showed a later onset of disease manifestations, a lower viremia and a delay in mortality when compared to mice infected with WT virus (Fig 1F, Supplementary table 1). This illustrated that production of sfRNAs also promotes replication and pathogenesis of ZIKV *in vivo*.

In summary, sfRNAs are required for optimal virus replication in vertebrate cells and *in vivo*, thereby increasing CPE and disease, respectively.

### Production of sfRNAs is required for ZIKV infection in the placenta and dissemination into foetal brain

To determine if production of sfRNAs contributes to ZIKV pathogenesis during pregnancy, an established animal pregnancy model^17, 18^ with mice deficient in the IFN-*α*/*β* receptor (IFNAR^-/-^) was used. IFNAR^-/-^ mice usually survive infection with Brazilian (Asian lineage) viruses, but die after infection with African lineage ZIKVs^16^. The xrRNA1’ and xrRNA2’ mutants of the Brazilian ZIKV strain Natal (ZIKV_Natal_) were thus constructed (Supplementary Figure 2A) and pregnant dams were inoculated with either WT or the sfRNA-deficient ZIKV_Natal_ viruses (10^4^ FFU per mouse) at E12.5. All three viruses were able to establish viremias in IFNAR^-/-^ dams (Fig 2A), with mutant viruses showing slightly lower titers than WT virus, although the differences did not reach statistical significance (Fig 2A, Supplementary Table 2). The mutant and WT viruses had no significant effect on placental or fetal weights (Supplementary Figure 2B) or on proportion of deformed fetuses (Fig. 2B, C). WT and xrRNA1’ ZIKV established productive infections in placenta, whereas no virus titers were detected in placentas of dams infected with the xrRNA2’ mutant (Fig 2B), despite similar viral titres in blood to those seen for the xrRNA1’ mutant. This indicated that production of sfRNAs is crucial for ZIKV infection of and replication in the placenta. Moreover, despite the presence of virus in placenta at levels comparable to WT, the xrRNA1’ mutant was detected in only one fetal head (3% infection rate, Fig 2C), whereas WT ZIKV was detected in 11 fetal heads (22% infection rate, Fig 2C). This indicated that production of sfRNA is crucial for dissemination of ZIKV into fetal heads. All fetal heads were negative in dams infected with the xrRNA2’ mutant (Fig 2C), as might be expected since placentas were also not infected (Fig 2B).

**Figure 2.**
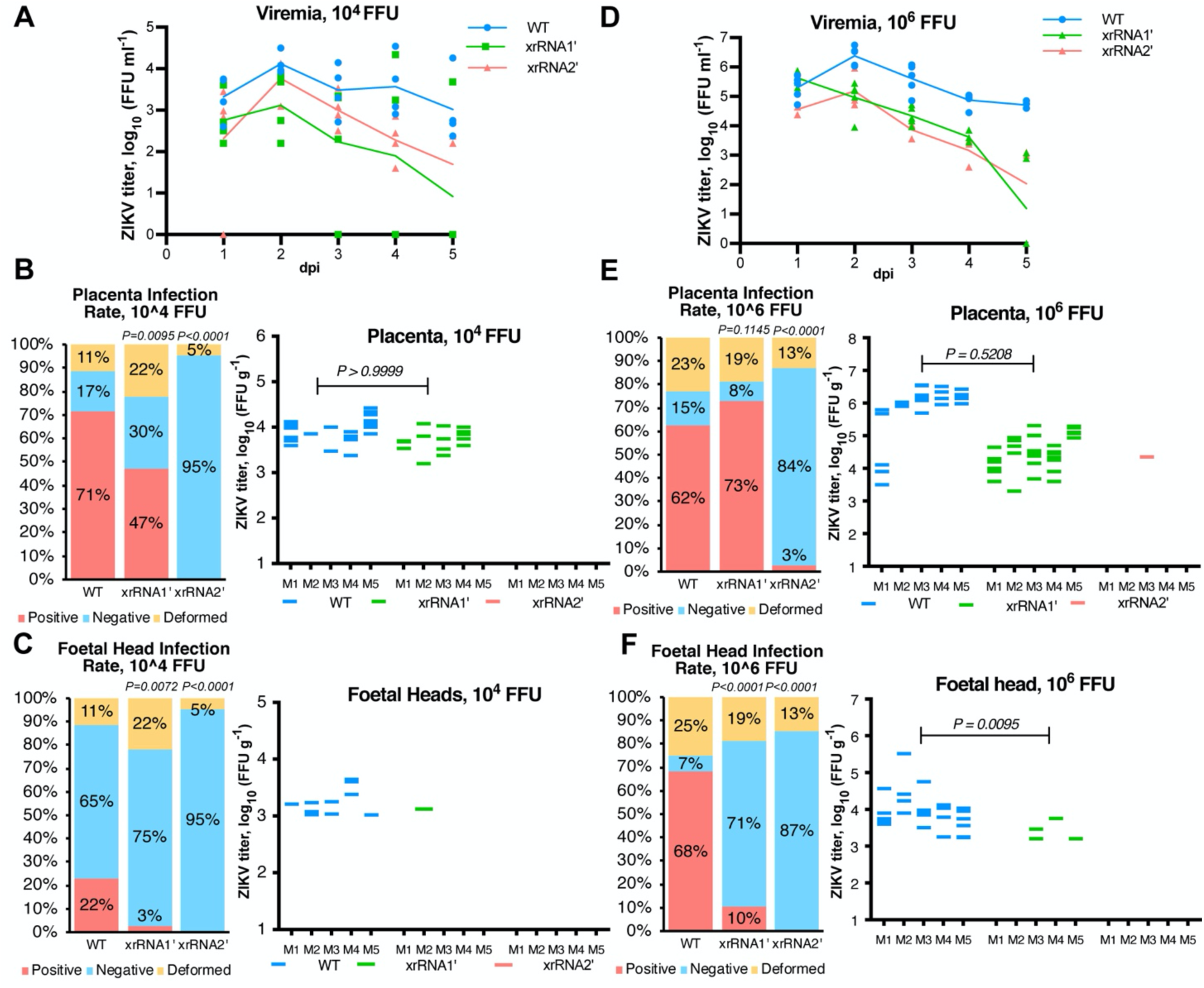
Production of ZIKV sfRNA is required for placental infection and dissemination of the virus into foetal heads in mouse pregnancy model. **(A, D)** Blood viremia in ZIKV-infected dams. Pregnant IFNAR -/- mice were inoculated with indicated doses of WT or sfRNA-deficient ZIKV Natal via subcutaneous injection. Blood samples were collected by tail bleeding. **(B, E)** Infection rate and viral titres in placenta samples collected from infected mice. **(C, F)** Infection rates and viral titres in mouse foetal heads. In (B, E, C, F) foetal material was collected at 5pdi, weighed, homogenised and used for virus titration by foci-forming assay. Viral titres were normalised to the tissue weight. The only virus-positive placenta sample in E for xrRNA2’ mutant contains reverse mutation to the WT sequence. Statistical analysis of infection rate is by Fisher’s exact test, P-values are in comparison to WT-infected group. Statistical comparison of virus titres was performed by Mann-Whitney U-test.

To assess the effects of a higher viral inoculation dose, dams were infected with a 100-times higher dose of each virus (10^6^ FFU). Viremia levels for all the viruses increased by ≈1-2 logs when compared to 10^4^ FFU infectious dose (Fig 2D, Supplementary figure 2C). Although again the levels for the WT virus were higher, the difference with the mutants was statistically significant only at 5 dpi (Supplementary table 3). While higher viremias resulted in increase in the placental infection rates for the xrRNA1’ mutant from 47% to 73%, the placental infection rate for the xrRNA2’ mutant virus only rose from 0% to 3% (Fig 2E). The latter represented a single placenta, with the follow up sequencing of the viral RNA identified reversion to the WT sequence (data not shown). The 22% dissemination into fetal heads for the WT virus (Fig. 2C) rose to 68% after infection with higher viral dose (Fig. 2F), and for the xrRNA1’ mutant this proportion increased from 3% to 10%. However, the infection rate for the xrRNA2’ mutant virus remained zero (Fig. 2F).

These results in pregnant animals clearly demonstrated that the sfRNAs of ZIKV are required for efficient viral replication in placenta and for transplacental migration and fetal brain infection.

### sfRNA facilitates ZIKV-induced apoptosis in neural progenitor cells leading to disintegration of human brain organoids

To gain insights into the effects of ZIKV sfRNAs on fetal neurovirulence and neuropathogenesis in humans, iPSC-derived human cerebral organoids (hCOs)^19^ were used as an *ex vivo* model of a developing human brain. Overt shrinkage of hCOs and loss of their typical structure by 6 days post infection (dpi) was evident for mutant and WT ZIKV_MR766_-infected organoids (Fig 3A), with reductions in organoid sizes compared to uninfected organoids clearly evident at later time points (Fig 3A). However, only WT virus infection caused CPE and complete disintegration of hCOs by 15-18 dpi, whereas organoids infected with either of the sfRNA-deficient mutants, although remaining smaller than uninfected organoids, survived through the course of the infection (Fig 3A). Viral titers early in infection with the WT virus (2-7 dpi) were higher than in infection with the mutant viruses (Fig 3B, Supplementary table 4). Consistent with these higher titers, on 3 dpi (Fig 3B), WT virus-infected organoids also had significantly more infected cells than organoids infected with the mutant viruses (Fig 3C, D).

**Figure 3.**
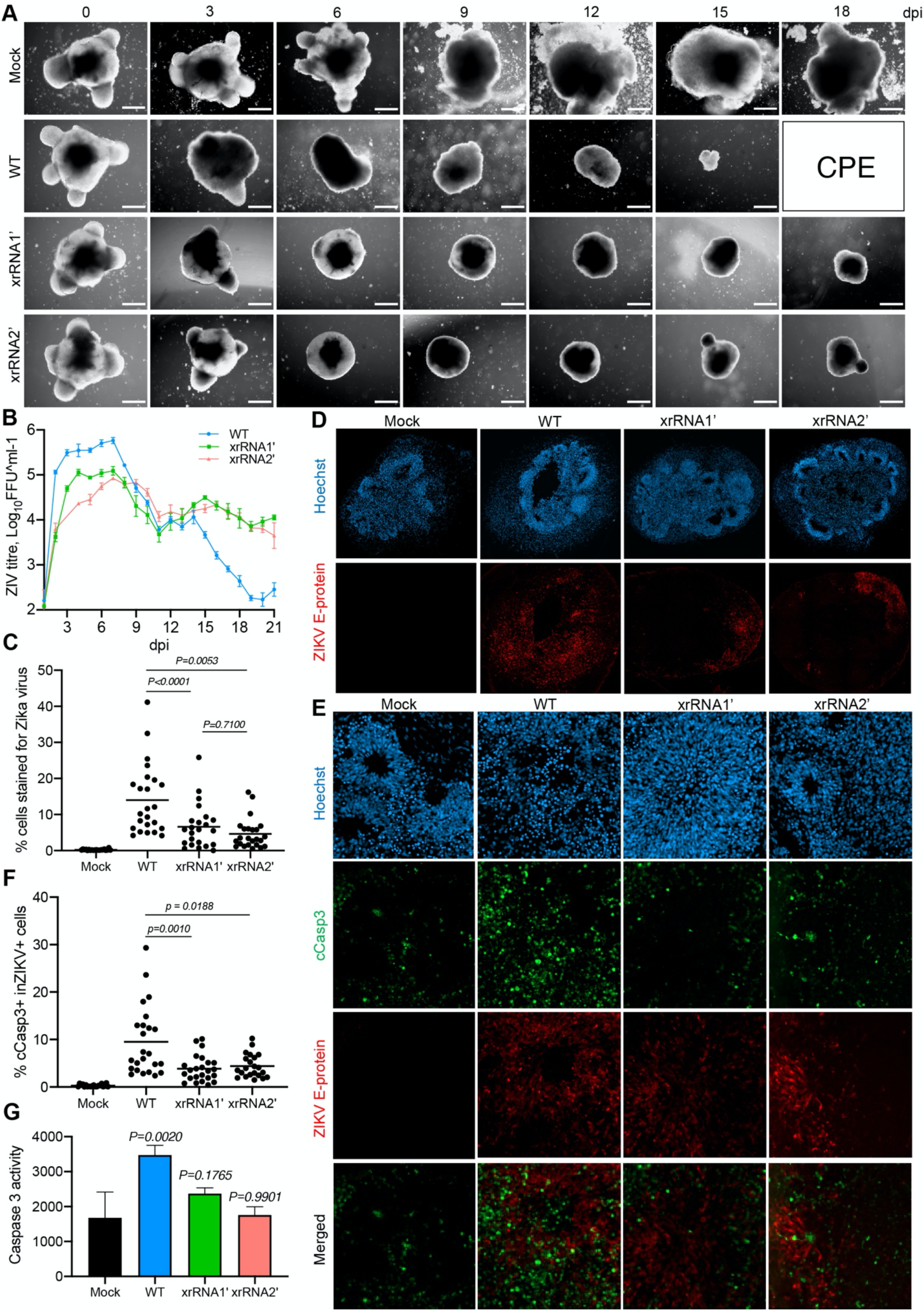
The sfRNA facilitates replication and cytopathogenicity of ZIKV in human brain organoids and promotes apoptosis of neural progenitors. **(A)** Morphology of human cerebral organoids infected with WT and sfRNA-deficient ZIKV. **(B)** Viral titres in culture fluids from human cerebral organoids infected with WT and sfRNA-deficient ZIKV shown in (A). The 14 days old organoids were infected with 10^4^ FFU ZIKV. Culture fluids were sampled daily and replaced with the equal volume of fresh growth medium. Viral titres were determined by foci-forming assay. Values are the means from three biologically independent organoids ±SD. Statistical analysis is summarised in Supplementary table 4. **(C, D)** Infection efficiency of WT and sfRNA-deficient ZIKV in human cerebral organoids. **(E, F)** Effect of sfRNA production on caspase-3 activation in ZIKV infected human brain organoids. In (C-F) Organoids were infected as in (B), then fixed and sectioned at 3dpi. ZIKV E-protein and cleaved caspase-3 were visualised by immunofluorescent staining and cell nuclei were stained with Hoechst fluorescent dye. Cells positive for ZIKV E-protein and cleaved caspase-3 (cCasp3) were counted across multiple sections of three biologically independent organoids. In (C, F) individual values for each section and mean values are shown. Statistical analysis was performed by Kruskal-Wallis test. Images in (A, D, E) are representative microphotograph of three biologically independent organoids that showed similar results. **(G)** Caspase 3/7 activity in immortalised human neural progenitor cells ReNCell infected with WT and sfRNA-deficient ZIKV. Cells were infected at MOI=1 and caspase 3/7 activity was assessed at 72hpi. Values are the means from three biological replicates ±SD. Statistical analysis was performed using one-way ANOVA, all comparisons were to mock.

Apoptosis of immature neurons and neural progenitors has been identified as one of the reasons for ZIKV-induced microcephaly^20^. Immunohistochemical detection of cleaved caspase-3 (cCasp3) was used to elucidate the effect of sfRNA on induction of apoptosis in hCOs. Sections of organoids fixed at 3 dpi with WT or the sfRNA-deficient ZIKV mutants demonstrated that in organoids infected with WT ZIKV a significantly higher proportion of infected cells were positive for cleaved Caspase 3, when compared with organoids infected with sfRNA-deficient mutant viruses (Fig 3E, F). In agreement with these data, analysis of caspase-3/7 activity in a homogeneous monolayer culture of human neural progenitor cell line ReNcell showed significant increase in casp-3/7 activity in response to infection with WT but not sfRNA-deficient mutant viruses at 72hpi (Fig 3G) even though cells infected with either virus had similar viral loads at this time point (Supplementary Fig 3). Therefore, observed increase in caspase3/7 activity can be attributed to the specific function of sfRNA in virus-induced apoptosis.

### ZIKV sfRNA inhibits signalling pathways that converge at STAT1

The data in Fig. 1F and Fig. 2B and E argue that sfRNAs facilitate ZIKV pathogenesis in AG129 mice deficient in receptors for IFN-*α*/*β*/*γ* and are required for infection of placenta in mice deficient in IFN-*α*/*β* receptor. This suggests that sfRNA may affect other, type I/II IFN response-independent, antiviral pathways. Transcriptome-wide gene expression profiling of cells infected with WT and sfRNA-deficient ZIKV was thus performed to fully map the effects of sfRNA on the entire spectrum of host responses. The human placental cell line BeWo was used for this experiment as it is known to support ZIKV replication^21^, is capable of responding to all three types of IFNs^21, 22^ and has a placental origin (and thus relevant to ZIKV tropism).

BeWo cells were infected with WT ZIKV or xrRNA2’-mutant viruses and at 72 hpi and total RNA was isolated and used for RNA-Seq. Notably, both viruses produced similar amount of intracellular viral RNA at this time point (Supplementary figure 4A). Principal component analysis of the library size-normalized RNA-Seq counts showed clear separation of samples infected with WT and xrRNA2’ ZIKV (Supplementary figure 4B). This indicates for the presence of gene expression component specifically associated with sfRNA production. Differential gene expression analysis demonstrated that multiple genes, including known interferon-stimulated genes (ISGs) such as Mx1, Mx2 and ISG15, showed stronger induction in response to xrRNA2’ mutant virus compared to WT virus (Fig 4A, Supplementary table 5). Subsequently, the gene ontology (GO) and KEGG pathway enrichment analyses identified type I IFN signaling as the most enriched biological process inhibited by sfRNA in these cells (Fig 4B). In addition, the enrichment analyses revealed that ZIKV sfRNAs inhibited IFN-*γ*, NF-*κ*B and TNF signaling (Fig 4B). While IFN-*γ* was not produced by these cells (GEO dataset GSE171648), STAT2 degradation by ZIKV^23^ is known to skew signaling towards the STAT1/STAT1 pathway^24^, which is the primary factor activated by in IFN-*γ* signaling. As a result, a prominent signature associated with stimulation of IFN-*γ* signaling emerges (Fig 4C). Moreover, sfRNAs appeared to decrease activation of “negative regulation of apoptotic processes” (Fig 4B, left panel) and slightly, albeit significantly induce p53-signalling (Fig 4B, right panel), which provides additional evidence for pro-apoptotic function of sfRNAs. Notably, the sfRNA-affected genes related to IFN responses and antiviral defense were represented by ISGs, receptors for IFN*α*/*β* and *γ*, and the components of IFN*α*/*β*/*γ*-signaling, whereas IFNs themselves were not identified as sfRNA-affected genes at this time point (Fig 4C). This suggests that ZIKV sfRNAs act downstream of IFN*α*/*β*/*γ* receptors, while not affecting IFN*α*/*β* production. The expression of ISGs was significantly higher in cells infected with xrRNA2’ ZIKV compared to WT ZIKV. A similar pattern was also observed in expression of pro-inflammatory (TNF-signaling pathway) and anti-apoptotic genes. Collectively, the results of transcriptome-wide gene expression profiling demonstrated that ZIKV sfRNAs inhibit type I and type II IFN signaling, TNF signaling and promote apoptosis. Potentially, they also inhibit type III IFN response, which shares gene expression signatures with the response to type I IFN and therefore cannot be distinguished from type I IFN pathway using current bioinformatics annotations.

**Figure 4.**
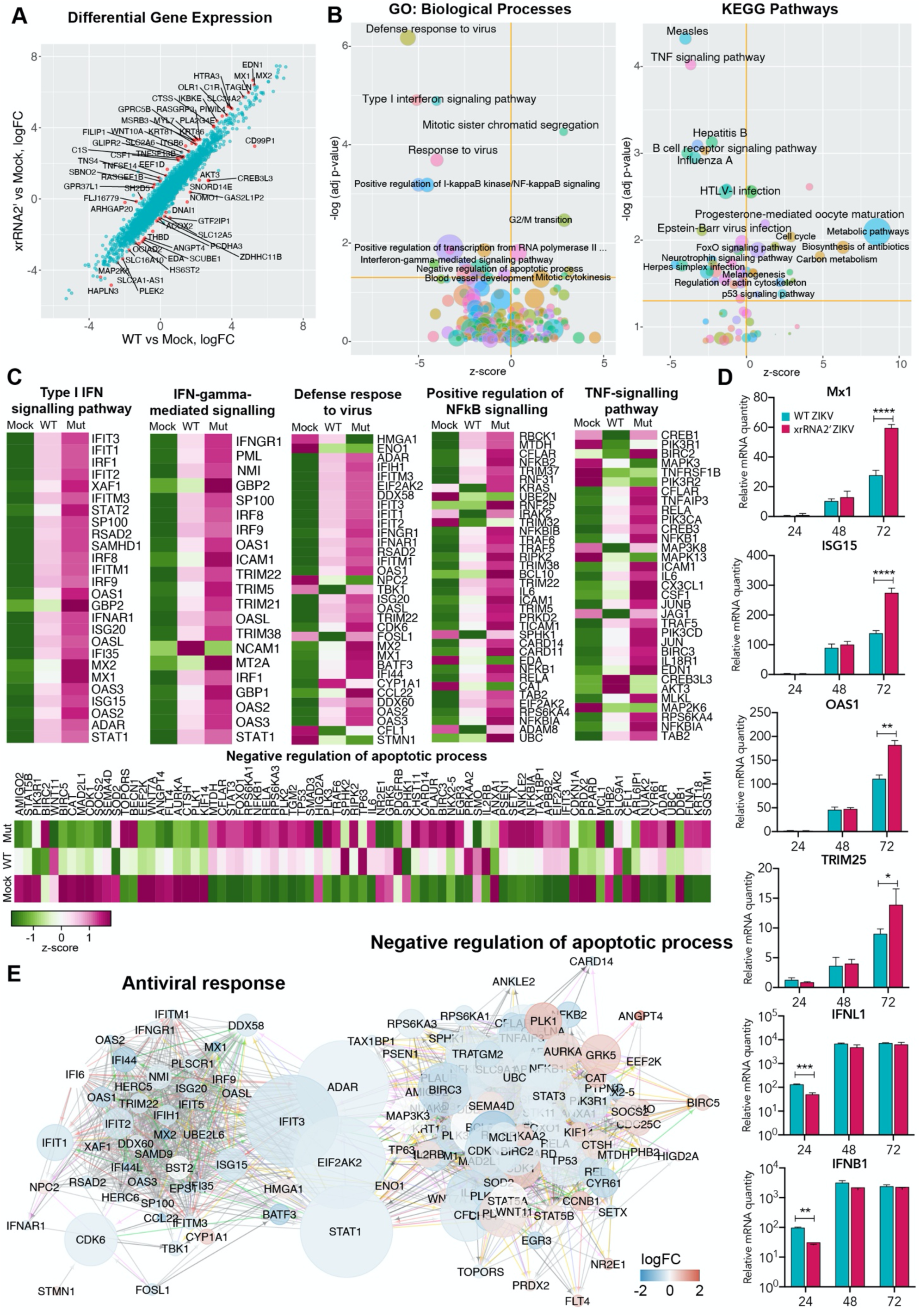
Effect of sfRNA production on expression of host genes in ZIKV-infected human placental cells. **(A)** Differential gene expression in BeWo cells infected with WT and xrRNA2’ ZIKV. Points indicating the genes that responded differently (FDR-adjusted P-value < 0.05) to infection with WT and mutant viruses are shown in red. **(B)** Pathways and biological processes affected by production of ZIKV sfRNA. The sfRNA-affected genes identified in (A) were subjected to GO and KEGG enrichment analysis. The data on enrichment significance was then combined with expression values for each gene and z-scores were calculated. Z-scores show directionality in expression changes of the genes associated with each sfRNA-affected process – negative indicate for inhibition and positive indicate for activation. **(C)** Expression of the individual genes associated with biological processes affected by production of ZIKV sfRNA in BeWo cells. Values are Z-scores with green indicating for negative and purple for positive and are the means from three biological replicates. **(D)** Expression of ISGs and IFNs in BeWo cells infected with WT virus (cyan bars) and xrRNA2’ mutant virus (magenta bars) ZIKV. Cells were infected at MOI=1, total RNA was isolated at the indicated time points and used for qRT-PCR. Relative mRNA quantity was determined using *ΔΔ*Ct method and is relative to Mock with normalisation to *GAPDH*. The values are the means of three biological replicates ±SD. Statistical analysis was performed using multiple t-tests; \**p<0.05*, \*\**p<0.01*, \*\*\**p<0.001*, \*\*\*\**p<0.0001*. **(E)** Network of interactions between the sfRNA-affected genes involved in antiviral response and apoptosis. Two individual networks were reconstructed form the genes identified in (C) using Genemania CytoScape plug-in and then merged. The resulted network was subjected to CytoScape network analysis to calculate the Betweennes Centrality values as a measure for the weight of each nod in the combined network. Size of the nods indicates for Betweenness Centrality, colour of the nods indicates the difference in gene expression between the cells infected with WT and xrRNA2’ mutant ZIKV. Positions of the nods are relative to their connectivity within and between the two subnetworks.

To validate the results of RNA-Seq we performed qRT-PCR for selected ISGs, and for IFN*β* and IFN*λ* in infected BeWo cells at different time points after infection. Consistent with the transcriptomics data, we observed inhibitory effect of sfRNAs on expression of ISGs later in infection (72 hpi), while the expression of IFN*β* and IFN*λ* was similar at this time point (Fig 4D). In addition, we found that replication of sfRNA-deficient mutant viruses is restored nearly to the levels of WT virus infection in IFNAR-deficient cells (Supplementary figure 5). These results further reinforce the notion that ZIKV sfRNAs inhibit IFN signaling but not IFN production.

To further dissect the molecular processes affected by ZIKV sfRNAs, we queried whether the regulatory networks associated with antiviral response and apoptosis intersect. To this end we determined the relative weight of each individual gene in these networks as an indication of their importance in the respective signaling. The network analysis identified STAT1, ADAR, IFIT3 and EIF2AK2 (PKR) as shared components of the antiviral response and negative regulation of apoptosis (Fig 4E). Moreover, these genes also have the biggest weight in the joint network as indicated by their “betweenness centrality” parameter (Fig 4E). The involvement of these genes in antiviral response and negative regulation of apoptosis suggests that their inhibition should simultaneously impair antiviral response and promote apoptosis. Therefore, we concluded that these genes may represent molecular targets of sfRNAs. Among these proteins, ADAR, IFIT3 and PKR are the effectors and the most downstream components of the type I and type III IFN signalling^25^ and were therefore deemed unlikely candidates for mediating the effect of sfRNA on expression of multiple ISGs. STAT1, however, is a key regulatory component in all types of IFN signaling. It is also known to antagonize apoptosis in cells exposed to the IFN-*γ*^26^. In addition, analysis of pathways differentially regulated by WT and sfRNA-deficient virus infections using Ingenuity Pathway Analysis software identified STAT1 as the most significant direct upstream regulator of the differentially expressed genes affected by sfRNA (Supplementary Table 6), indicating the key role of sfRNA in inhibiting STAT1-regulated pathways. Based on these results we hypothesized that STAT1 represents the key target of sfRNAs, which allows them to modulate multiple host response pathways.

### ZIKV sfRNA inhibits phosphorylation and nuclear translocation of STAT1 in infected cells

To elucidate the effect of sfRNAs on STAT1 activation, the levels of total and phosphorylated STAT1 were determined in Vero cells infected with WT or xrRNA2’ ZIKV after treatment with IFN-*α*. We found that IFN-*α* induced substantial accumulation of Tyr701- phosphorylated STAT1 (pSTAT1) in mock-infected cells and cells infected with the sfRNA-deficient ZIKV, but not in cells infected with the WT virus (Fig 5A). Equal amounts of viral NS3 protein were detected in both infections, indicating that sfRNA deficiency did not significantly affect viral loads in these cells. The total STAT1 levels were not affected by the infections with both viruses (Fig 5A), indicating that sfRNAs inhibited STAT1 phosphorylation rather than affecting STAT1 expression. In addition, the immunofluorescent microscopy revealed that infection with WT, but not with xrRNA2’ ZIKV, efficiently prevented accumulation of pSTAT1 in the nuclei of infected cells (Fig 5B), which suggests that transcription of STAT1-regulated ISGs would be impaired by sfRNAs.

**Figure 5.**
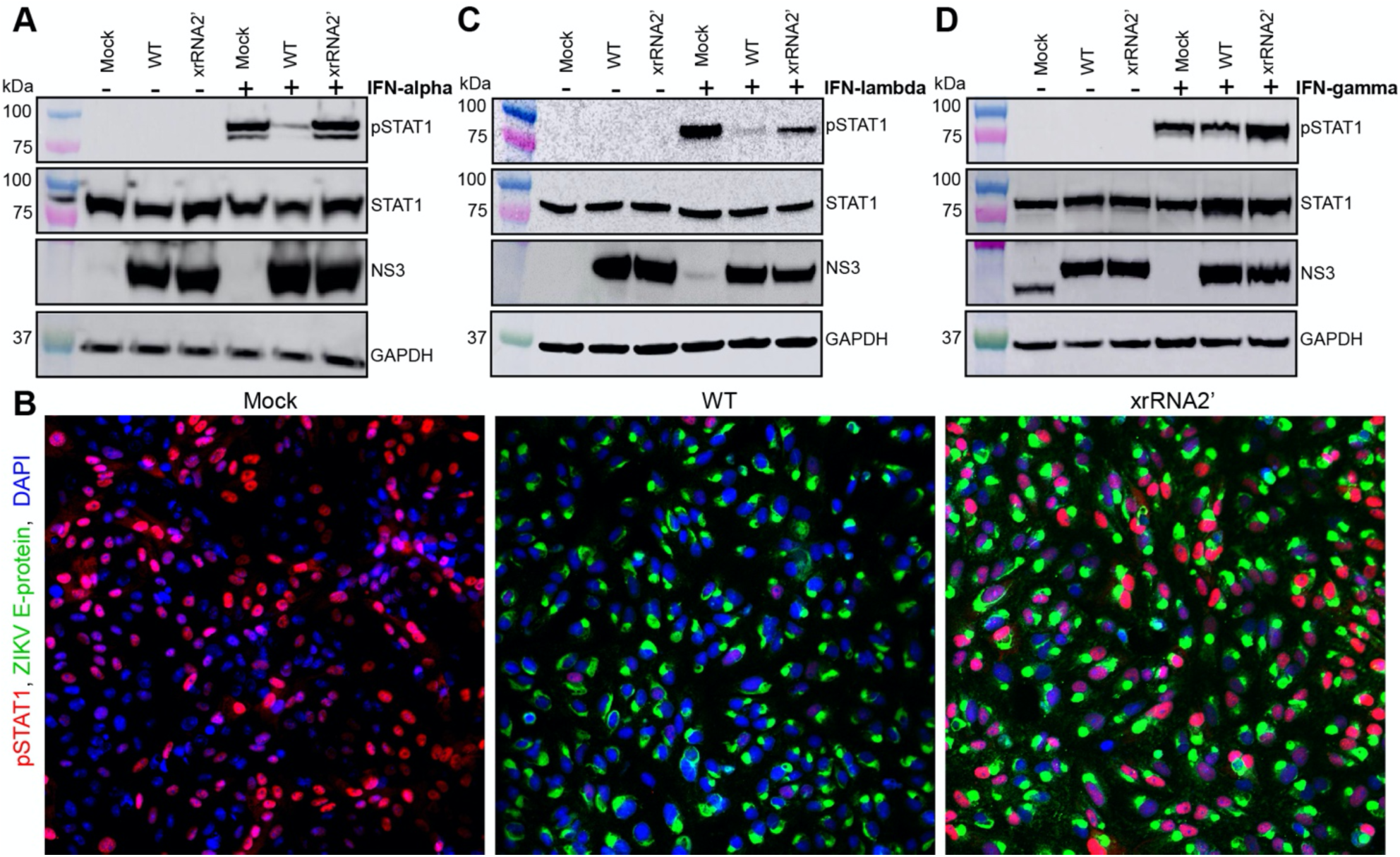
ZIKV sfRNA inhibits type I, II and III IFN signalling by supressing phosphorylation and nuclear translocation of STAT1. **(A, C, D)** Effect of ZIKV sfRNA on phosphorylation of STAT1 in response to type I (A), type III (C) and type II (D) IFN. Vero (A, D) or HTR-8 cells (C) were infected with WT or xrRNA2’ ZIKV or left uninfected (Mock). At 48hpi cells were treated with human IFN*α*1 (A), *λ*1 (C) or *γ* for 20 (A, D) or 60 (C) minutes. Levels of Tyr701-phosphorylated STAT1 (pSTAT1) and total STAT1 indicate for phosphorylation and expression of STAT1, respectively. Levels of ZIKV NS3 indicate for viral loads in the infected cells. GAPDH levels indicate for the total protein input. (B) Immunofluorescent detection of Tyr701-phosphorylated STAT1 in ZIKV-infected Vero cells treated with IFN*α*1. Treatment and infection were performed as in (A), the image is the representative of three independent experiments that showed similar results.

We next tested the ability of sfRNAs to inhibit STAT1 phosphorylation in response to IFN-*λ* in human placental HTR-8 cells that have functional IFN-*λ* signalling^21^. The results demonstrated that sfRNAs reduced IFN-*λ*-induced phosphorylation of STAT1, while having no effect on the total STAT1 protein levels (Fig 5C). ZIKV sfRNA thus also disrupts type III IFN signaling by inhibiting phosphorylation of STAT1. Type II IFN signaling is exclusively regulated by STAT1, hence the effect of sfRNA on IFN-*γ* signaling was also examined. Vero cells that are capable of responding to IFN-*γ*^27^ were used for this experiment. The results showed less phosphorylated STAT1 in IFN-*γ*-treated cells infected with WT virus than in cells infected with xrRNA2’ mutant virus (Fig 5D), illustrating inhibition of IFN-*γ* signaling by sfRNA. Hence, we demonstrated that sfRNAs play a key role in inhibition of STAT1 phosphorylation in response to all three types of IFNs.

### ZIKV sfRNA interacts with viral protein NS5 to inhibit STAT1 phosphorylation

To further characterize the inhibitory activity of ZIKV sfRNAs on STAT1 phosphorylation and nuclear translocation, Vero cells were transfected with *in vitro-* generated ZIKV sfRNA or GFP RNA fragment (control), followed by treatment with IFN*α* (Supplementary figure 6). The sfRNA-transfected cells showed substantial accumulation of pSTAT1 in the nuclei, similar to the levels seen in cells transfected with control RNA and non-transfected cells (Supplementary figure 6B). Thus, sfRNAs alone do not inhibit nuclear translocation of pSTAT1, suggesting other viral or virus-induced host factors are required for this activity.

The activities of sfRNA in flaviviruses have previously been associated with interactions between sfRNA and host proteins^7, 8, 28^. These interactions are usually identified by an RNA affinity pull-down experiment in which lysates of uninfected cells are used with *in vitro* transcribed sfRNA ^29, 30^. However, given the lack of inhibitory activity of sfRNA alone on STAT1 phosphorylation and nuclear translocation in uninfected cells (Supplementary figure 6B), functional interactions of sfRNAs affecting STAT1 phosphorylation need to be studied in the content of virus infection. Since *in vitro* transcribed sfRNA used as a bait in pull-downs would compete with sfRNAs produced in cells infected with the WT virus, lysates of cells infected with the sfRNA-deficient virus were used for sfRNA affinity pull down assays. Streptavidin-binding aptamer-tagged ZIKV sfRNA and control GFP RNA (Supplementary figure 7) were generated by *in vitro* transcription and used in RNA affinity pull downs with lysate of cells infected with xrRNA2’ mutant virus to identify sfRNA-interacting proteins. Label-free quantitative mass spectrometry (SWATH-MS) was then used to identify proteins enriched in sfRNA pull-downs, compared to GFP RNA pull-down. This approach revealed ZIKV NS5 as the most significantly enriched sfRNA-interacting protein (Fig 6A). We therefore hypothesized that sfRNA is the viral factor that acts in conjunction with NS5 protein to provide efficient inhibition of STAT1 phosphorylation. NS5 of WNV^31^ and JEV^32^ are already known to inhibit STAT1 phosphorylation. However, NS5 ectopically expressed from plasmids was unable to inhibit STAT1 phosphorylation to the same extent as NS5 produced during infection, suggesting the requirement of another viral factor for this activity^33^.

**Figure 6.**
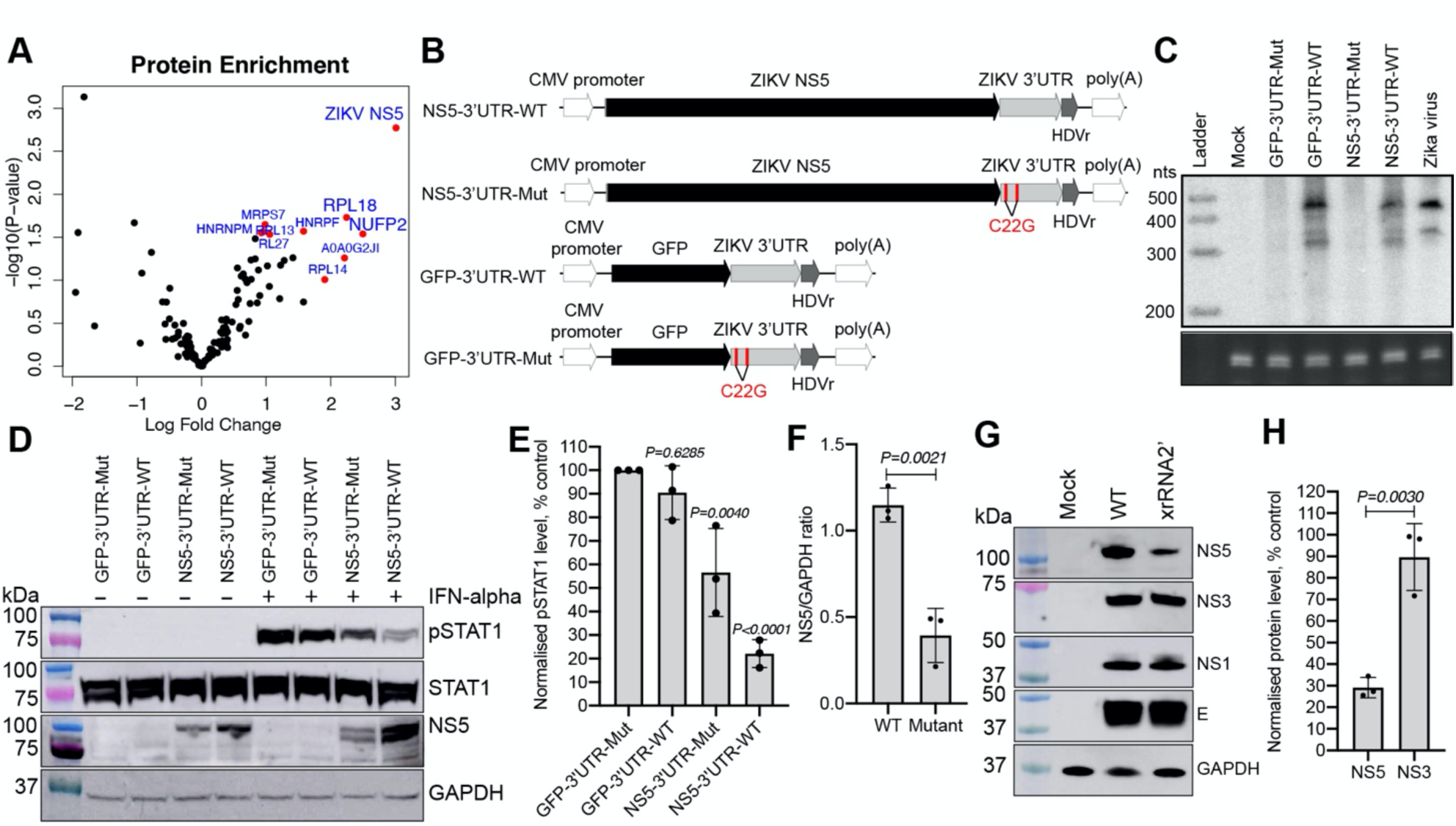
ZIKV sfRNA inhibits STAT1 phosphorylation by stabilising NS5 via direct binding. **(A)** The sfRNA-binding proteins identified in lysates of A549 cells infected with sfRNA-deficient ZIKV mutant using RNA affinity pull-down. Top 10 most significant sfRNA-interacting proteins are highlighted in red circles and named. The values are the means from three independent experiments. **(B)** Schematics of the reporter constructs generated for production of ZIKV sfRNA and NS5 alone and in combination. The C22G mutations that abolish XRN-1 resistance in each of ZIKV xrRNA elements and disrupt production of sfRNA are highlighted. **(C)** Northern blotting demonstrating production of sfRNA in HEK293T cells transfected with expression constructs shown in (B). RNA was isolated at 48h post transfection and 5ug was loaded per lane. Bottom panel shows 5.8S rRNA on an Et-Br-stained gel as a loading control. **(D)** Western-blot detection of phosphorylated STAT1 (pSTAT1), total STAT1 and ZIKV NS5 in HEK293T cells transfected with expression constructs shown in (B) and treated with IFN*α*. IFN-treatment was performed at 48hpt and cells were lysed for Western blotting after 15 min of treatment. GAPDH is shown as a loading control. **(E)** Quantification of pSTAT1 levels in (D) by densitometry. Normalised levels represent the ratios of pSTAT1 band density to GAPDH band density and expressed as a percentage relative to the values observed in the samples transfected with pcDNA-GFP-3’UTR-Mut plasmid. **(F)** Quantification of NS5 levels in (D) by densitometry. **(G)** Western-blot detection of structural (E-protein) and non-structural (NS5, NS3, NS1) proteins in the lysates of Vero cells infected with WT and sfRNA-deficient (xrRNA2’) ZIKV. Cells were infected at MOI=5 and assays were performed at 48hpi. **(H)** Quantification of the viral proteins levels in (G) by densitometry. Normalised levels represent the ratios of each protein band density to GAPDH band density and expressed as a percentage of the protein level in xrRNA2’-infected cells to WT ZIKV-infected cells. The blots in (C, D, G) are representative from three independent experiments that showed similar results. The values in (E, F, H) are the means from three independent experiments ±SD. Statistical analysis was performed by one-way ANOVA with Dunnett correction (E) or Student’s t-test (F, H).

To test this hypothesis, expression plasmids producing ZIKV NS5 and sfRNA (NS5-3’UTR-WT), ZIKV NS5 alone (NS5-3’UTR-Mut), GFP and ZIKV sfRNA (GFP-3’UTR-WT) and GFP alone (GFP-3’UTR-Mut) were generated (Fig 6B-D). HEK293T cells were transfected with each plasmid, treated with IFN*α* and analyzed by Western blot for pSTAT1 (Fig 6D). Consistent with the results of sfRNA transfection (Supplementary figure 6), similarly robust levels of pSTAT1 were observed in cells expressing only sfRNA (GFP-3’UTR WT) and cells transfected with a control construct producing neither sfRNA nor NS5 (GFP-3’UTR-mut) (Fig 6D,E). Cells that expressed NS5 without sfRNA (NS5-3’UTR-Mut) exhibited slightly decreased accumulation of pSTAT1 (Fig 6D,E), which illustrated that NS5 alone is capable of inhibiting STAT1 phosphorylation, but only to a limited extent. In contrast, accumulation of pSTAT1 was dramatically reduced in cells transfected with the construct expressing both ZIKV NS5 and sfRNA (NS5-3’UTR-WT), (Fig 6D,E). These results show that cooperative action of NS5 and sfRNA is required to efficiently inhibit STAT1 phosphorylation.

### ZIKV sfRNA stabilizes NS5 protein

Together with analysis of phosphorylated and total STAT1, we also assessed intracellular levels of NS5. Notably, cells co-expressing NS5 and ZIKV sfRNA had significantly higher levels of NS5 compared to cells expressing NS5 alone (Fig 6D, F). As cells were transfected with the same amounts of plasmid DNA, and the only difference between the control and sfRNA expressing plasmids was a mutation in two nucleotides of ZIKV 3’UTR that results in sfRNA-deficiency, we concluded that sfRNA is required for accumulation of NS5. Taken together with data in Fig 6A, these results indicate that sfRNA binds to and stabilizes NS5 protein.

To provide additional evidence that ZIKV sfRNA stabilizes NS5, the levels of non-structural (NS5, NS3, NS1) and structural (E) ZIKV proteins were assessed in Vero cells infected with WT and sfRNA-deficient ZIKV mutant xrRNA2’. Cells infected with the sfRNA-deficient ZIKV contained less NS5 than cells infected with the WT virus (Fig 6G, H). Infection with both viruses produced comparable levels of other viral proteins (Fig 6G, H). These results show that sfRNA stabilizes NS5 during viral infection and that the effect is specific for NS5.

ZIKV NS5 was previously shown to induce STAT2 degradation^23^ and a recent study demonstrated that sfRNA-deficient ZIKV mutant was also capable of this activity^34^, thereby excluding a role for ZIKV sfRNA in STAT2 degradation. Consistent with these results, we also showed that deficiency in sfRNA production and associated decrease in NS5 level did not affect STAT2 degradation in cells either infected with viruses or transfected with expression constructs (Supplementary figure 8). This suggest that STAT2 degradation can be efficiently induced by relatively small amount of NS5, while inhibition of STAT1 phosphorylation requires higher amounts of NS5 achieved by NS5 interaction with and stabilization by sfRNA.

## Discussion

Since our discovery of sfRNA^13^, it has attracted significant research interest^9^. Herein we demonstrate that sfRNA is required for efficient replication of ZIKV and that deficiency in production of sfRNA results in significant attenuation of virus replication in human cells and in a mouse model *in vivo*. Importantly, we found that deficiency in sfRNA prevented virus from infecting mouse placenta and disseminating into the fetal brain. We also demonstrated that ZIKV sfRNA promotes virus-induced CPE in cultured cells and apoptosis of neural progenitors in the infected developing human brain tissue. We found that sfRNA exerts these functions by inhibiting phosphorylation of STAT1 enabled by sfRNA binding to and stabilizing viral IFN antagonist, NS5 protein. This provides fundamental understanding of the role of sfRNA in subversion of host responses during ZIKV infection.

Mammalian cells produce three types of IFNs, of which type I IFN signaling is the most ubiquitous antiviral mechanism active against flaviviruses^35^. Type III IFNs are also involved in defense against flaviviruses, but function in tissue-specific manner. In particular, IFN-*λ* pathway is active in placenta and the female reproductive tract and plays a crucial role in protecting these tissues from ZIKV infection^21, 36^. Type II IFN is produced by NK cells as part of the innate immune response and subsequently by T cells as adaptive immunity develops^37^. Many types of cells, including placental trophoblasts^38^, express IFN-*γ* receptors and mount antiviral responses after exposure to IFN-*γ*^39^. To date, IFN-*γ* has been demonstrated to act against WNV and DENV to prevent systemic virus spread^40, 41^. In addition, treatment with IFN-*γ* was shown to reduce ZIKV replication in a human cell line^42^ and in primary human cells^43, 44^. Here, we showed that ZIKV sfRNAs play significant role in inhibiting the responses to IFN-*γ*, IFN-*α* and IFN-*λ* by preventing phosphorylation of STAT1.

Signal transduction from type I and type III IFNs relies on Tyr701-phosphorylation of STAT1, which then associates with phosphorylated STAT2 and IRF9 to form a transcription complex ISGF3, which translocates into the nucleus and up-regulates expression of ISGs^45^. In addition, pSTAT1 can form homodimers that mediate signal transduction from type II IFN and act as non-canonical signaling pathway in response to type I IFN^46^. Moreover, STAT1 plays a rather complex role in regulation of apoptosis, being able to exert pro- as well as anti-apoptotic functions depending on type of phosphorylation and cellular context^47^. In particular, STAT1 protects cells exposed to IFN-*γ* from apoptosis by activating transcription of genes that encode suppressors of apoptosis^26^. Here using transcriptomic data analysis of human placental cells infected with WT and sfRNA-deficient viruses we identified STAT1 as a common link between major antiviral pathways affected by sfRNA - IFN signaling and negative regulation of apoptosis.

Inhibition of STAT1 phosphorylation in flavivirus-infected cells was previously linked to the activity of viral protein NS5^31^. However, this activity was shown to be significantly lower for ectopically expressed NS5 in the absence of infection compared to that in infected cells, suggesting a requirement for another viral factor^33^. Here we identified sfRNA as this viral factor by showing that it binds to and stabilizes NS5, which results in NS5 accumulation to the levels required for efficient inhibition of STAT1 phosphorylation. To the best of our knowledge this is the first report demonstrating cooperation between noncoding viral RNA and a viral protein in inhibition of host antiviral responses. ZIKV NS5 is also known to trigger proteolytic cleavage of STAT2^23^. Consistent with a recent report^34^, we demonstrate that sfRNA is not required for this activity and that the small amounts of NS5 present in cells infected with sfRNA-deficient virus was sufficient to degrade STAT2.

The concentration-dependent requirement for NS5 in inhibition of STAT1 phosphorylation and concentration-independent requirement for NS5 in STAT2 degradation can be mechanistically explained based on our data and existing knowledge (Supplementary figure 9). Suppression of STAT1 phosphorylation by ZIKV NS5 was shown to require direct binding of NS5 to Hsp90, which inactivates HSP90^33^. Hsp90 is required for folding of Tyk2, the upstream kinase that phosphorylates STAT1. Inactivation of Hsp90 results in incorrect folding and subsequent proteasomal degradation of Tyk2, thereby preventing downstream STAT1 phosphorylation in response to IFN receptor signaling ^33^. Assuming NS5-Hsp90 binding is a reversible reaction, high levels of NS5 would be required to suppress activity of abundantly expressed Hsp90. Stabilization of NS5 by sfRNA results in accumulation of substantial amounts of NS5 which is required for inhibition of STAT1 phosphorylation. In contrast, the effect of NS5 on STAT2 is irreversible as it involves triggering the energy-dependent ubiquitination of STAT2, followed by its proteasomal degradation with NS5 being recycled. NS5 thereby acts as a catalyst^23, 48^ with small amounts of NS5 sufficient for STAT2 degradation.

Previous study indicated that NS5-induced STAT2 degradation switches signaling from STAT1/STAT2-driven responses to STAT1/STAT1-driven responses^24^ in ZIKV-infected cells. Herein, we propose that ZIKV counteracts type I and type III IFN responses by utilizing a small amount of available NS5 to induce STAT2 degradation and prevent formation of STAT1/STAT2 heterodimers. Removal of STAT2 switches signaling to the STAT1/STAT1 homodimer pathway which is used primarily by type II IFN response. sfRNA inhibits this pathway by binding to and stabilizing NS5 which enables NS5 to counteract STAT1 phosphorylation. Thus, the sfRNA-mediated inhibition of all 3 types of IFN response allows ZIKV to spread systemically, reach and replicate in placenta and eventually reach fetal brain where it induces apoptosis and neuropathogenicity.

## Methods

### Cells

Female African green monkey (*Cercopithecus aethiops*) kidney fibroblasts cells (Vero, ATCC – CCL-81), male human alveolar carcinoma cells A549 (ATCC – CCL-185), male human placental trophoblast cells BeWo (ATCC – CCL-98), human placental trophoblast cells HTR-8 (ATCC – CRL-3271), human embryonic kidney cells HEK-293T (ATCC – CRL-3216) and *Aedes Albopictus* larvae cells C6/36 (ATCC – CRL-1660) were obtained from ATCC. Wild type and IFNAR^-/-^ mouse embryonic fibroblasts (MEF) have been generated previously^5^. ReNcell Human Neural Progenitor Cells (hNPCs) were provided by Professor Bryan Day and Dr. Ulrich Baumgartner from QIMRB, QLD, Australia. Vero, MEF, HEK-293T and HTR-8 cells were cultured in Dulbecco’s modified Eagle’s medium (DMEM) supplemented with 10% (v/v) fetal bovine serum (FBS). A459 and BeWo cells were maintained in Ham’s F-12K (Kaighn’s) medium supplemented with 10% (v/v) FBS. ReNcell were cultured KnockOut^TM^ DMEM/F-12 supplemented with 2% StemPro^TM^ Neural Supplement, 10 μg of EGF Recombinant Human Protein, 5 μg of FGFb Recombinant Human Protein and 1% Gluta-Max^TM^ (Gibco) in flasks coated with Matrigel Matrix Basement Membrane (Corning; 1:100 dilution in PBS). C6/36 cells were cultured in Roswell Park Memorial Institute 1640 medium (RPMI 1640) supplemented with 10% (v/v) FBS and 10 mM HEPES pH7.4. All vertebrate cells were incubated at 37 °C with 5% CO2. Insect cells C6/36 were cultured at 28 °C in sealed containers. All cell culture media and reagents were from Gibco, USA unless specified otherwise.

### Viruses and infection of cells

Zika virus strains MR766 was obtained from Victorian Infectious Diseases Reference Laboratory. Natal strain of ZIKV was previously assembled from synthetic DNA fragments^49^. The viruses were passaged once in C6/36 cells and viral titres were determined by foci forming assay on Vero76 cells. The viral genomes were sequenced and confirmed to match GeneBank reference sequences MK105975 for ZIKV_MR766_ and KU955594 for ZIKV_Natal_. All infections were performed at the indicated Multiplicity of Infection (MOI) by incubation of cells with 50 μl of inoculum per cm^2^ of growth area for 1h at 37 °C. Inoculated cells were then maintained in the growth medium containing a reduced amount of FBS (2%) to prevent overgrowth.

### RNA isolation

Viral RNA was isolated from cell culture fluids using the NucleoSpin RNA Virus Kit (Macherey-Nagel, Germany). Total RNA from cells was isolated using TRIreagent (Sigma, USA). RNA from pull-down samples was isolated using TRIreagent LS (Sigma, USA). All RNA isolation procedures were conducted according to the manufacturer’s instructions.

### DNA isolation

DNA extraction from agarose gels was performed using Monarch DNA Gel Extraction Kit (NEB, USA). DNA isolation from reaction mixtures was performed using Monarch PCR & DNA Clean-up Kit (NEB, USA). Plasmid miniprep DNA isolation was performed using Wizard Plus SV Minipreps DNA Purification System (Promega, USA). Plasmid maxiprep DNA isolation was performed using NucleoBond Extra Maxi Plus EF Kit (Macherey-Nagel, Germany).

### Generation of sfRNA-deficient ZIKV mutants

The sfRNA-deficient mutants of ZIKV_MR766_ xrRNA1’ and xrRNA2’ were generated previously^14^. To generate xrRNA1’ and xrRNA2’ mutants of ZIKV_Natal_, the pUC19 plasmid containing the fragment of ZIKV cDNA with full-length 3’UTR ^50^ was used a template for PCR-directed site-specific mutagenesis. The primers ZIKV_xrRNA1’F and ZIKV_xrRNA1’R or ZIKV-xrRNA2’F and xrRNA2’R (Supplementary table 7) were used to introduce C to G substitution into the position of each xrRNAs critical for XRN-1 resistance ^4, 14^. Mutagenesis was performed using Q5 Site-Directed Mutagenesis Kit (NEB, USA) according to the manufacturer’s instructions. The cycling conditions for PCR were as follows: 98 °C – 30 s; 25 x (98 °C – 10 s, 55 °C – 30 s, 72 °C – 45 s); 72 °C – 2 min. Mutagenized constructs were transformed into NEB5*α* highly competent cells (NEB, USA), plasmids were isolated, and presence of mutations was confirmed by Sanger Sequencing. Fragments containing the mutations were then amplified from the plasmid using primers Natal 7F and Natal 7R (Supplementary table 7) and used for the assembly of infectious cDNA via circular polymerase extension reaction (CPER)^49^.

To generate other DNA fragments for CPER, viral RNA was isolated from culture fluids of C6/36 cells infected with ZIKV_Natal_ at 5 dpi and used as a template for first strand cDNA synthesis with SuperScript IV reverse transcriptase (Invitrogen, USA). Each RT reaction contained 11 µl of viral RNA, and 2 pmol of the reverse PCR primer for the corresponding fragment. RNA was denatured for 5 min at 95 °C and annealed at 65 °C followed by cDNA synthesis at 55 °C for 1h. RNA was then removed by incubation with 1 µl of RNase H (NEB, USA) for 20 minutes at 37 °C. RNase H-treated cDNA was used as a template for PCR with PrimeStar GXL Polymerase (Takara, Japan) and primers listed in Supplementary table 7. The cycling conditions were 3 min at 98° C; 40 cycles of 10 s at 98 °C, 15 s at 55 °C, 4 min at 68 °C; and a final extension for 5 min at 68 °C. PCR products were then separated in 1% agarose gel and DNA was gel-purified.

Assembly of the infectious cDNA was conducted using the circular polymerase extension reaction (CPER)^51, 52^. PCR-amplified cDNA fragments were mixed with the UTR- linker fragment containing OpIE promoter and either WT ZIKV_Natal_ 3’UTR fragment or one of the two mutated fragments. CPER mixtures contained 0.1 pmol of each DNA fragment and 2 µl of PrimeStar GXL DNA polymerase (Takara, Japan) in a total reaction volume of 50 µl. The cycling conditions were 2 min at 98 °C; 20 cycles of 10 s at 98 °C, 15 s at 55 °C, 12 min at 68 °C; and a final extension for 12 min at 68 °C. The resulted PCR products were transfected directly into C6/36 cells using Lipofectamine 2000 (Invitrogen, USA). Briefly, each CPER product was diluted in 100 µl of Opti-MEM (Invitrogen, USA) and mixed with of 12.5 µl of Lipofectamine 2000 dissolved in 150 µl of Opti-MEM. DNA-lipid complexes were incubated for 5 min at room temperature and added onto C6/36 cells grown to 80% confluence in the wells of 6-well plates. At 24h after transfection, cell culture medium was replaced and cell culture supernatant containing passage (P0) viruses were harvested at 5 dpi. Virus titres were determined by IPA and P0 viruses were used to infect C6/36 cells at an MOI of 0.1 to produce high titre P1 virus stocks.

### RT-PCR

To confirm retention of mutations in P0 and P1 viruses, viral RNA was isolated from virus samples and ZIKV 3’UTR was PCR amplified using primers Natal 3’UTR F and Natal 3’UTR R. PCR products were gel-purified and analysed by Sanger Sequencing. RT-PCR was performed using SuperScript III One-Step RT-PCR Kit with Platinum Taq (Invitrogen, USA) according to the manufacturers’ recommendations. 25 µl reaction mixtures containing 5 µl of viral RNA and 5 pmol of each primer were incubated under the following cycling conditions: 60 °C – 15 min, 94 °C – 2 min followed by 30 cycles of 94 °C – 15 sec, 55 °C – 30 sec, 68 °C – 1 min with final extension at 68 °C for 5min.

### Foci-forming immunoassay

Viral titers were determined by foci-forming assay. Ten-fold serial dilutions of cell culture fluids mouse serum or tissue homogenates were prepared in DMEM media supplemented with 2% FBS and 25 µl of each dilution were used to infect 10^5^ C6/36 cells pre-seeded in 96 well plates. After 1 h of incubation at 28 °C with the inoculum, 175 µl overlay media was added to each well. The overlay media contain a volume of 2x M199 medium (supplemented with 5% FCS, 100 μg/ml streptomycin, 100 U/ml penicillin, and 2.2 g/L NaHCO3) and another volume of 2% carboxymethyl cellulose (Sigma-Aldrich, USA). At 3 days-post infection, overlay medium was removed and cells were fixed with 100 µl/well of cold 80% acetone (diluted in 1x PBS) for 30 min at -20°C. Then, after fixative solution was removed and the monolayer was fully dried, plates were blocked for 60 min with 150 µl/well of ClearMilk blocking solution (Pierce, USA), and incubated with 50 μl/well of 4G2 mouse monoclonal antibody to ZIKV envelope protein diluted 1:100 for 1h, followed by 1h incubation with 50 μl/well of 1:1000 dilution of goat anti-mouse IRDye 800CW secondary antibody (LI-COR, USA). All antibodies were diluted with Clear Milk blocking buffer (Pierce, USA) and incubations were performed at 37 °C for 1h. After each incubation with antibody, plates were washed 5 times with phosphate buffered saline containing 0.05% Tween 20 (PBST). Plates were then scanned using an Odyssey CLx Imaging System (LI-COR) using the following settings: channel = 800 and 700, intensity = auto, resolution= 42 μm, quality= medium and focus= 3.0 mm.

### Plaque assay

Ten-fold serial dilutions of virus samples were prepared and 200 ul of each was used to inoculate Vero cells grown in 6-well plates by incubation for 1 h. Cells were then overlayed with DMEM containing 0.5% low melting point agarose (Rio-Rad, USA) and 2% FBS. At 72 h post-infection cells were fixed with 4% PBS-buffered formaldehyde for 1h, stained with 0.1% crystal violet for 30min and washed with water.

### Northern Blotting

Detection of sfRNA was performed by Northern blotting ^53, 54^. Total RNA (10ug) was mixed with an equal volume of Loading Buffer II (Ambion, USA), denatured by heating at 85 °C for 5 min and then placed on ice for 2 min. Denatured RNA was used for electrophoresis in 6% polyacrylamide TBE-Urea gels (Invitrogen, USA), which was performed at 180V for 90 min in 1x Tris-Borate-EDTA buffer pH8.0 (TBE). Gels were stained with ethidium bromide to visualise rRNA and documented using GelLogic 212PRO imager (Carestream, Canada). RNA was then electroblotted onto Amersham Hybond-N^+^ nylon membrane (GE Healthcare, USA) for 90 min at 35V in 0.5x TBE using the TransBlot Mini transfer module (Bio-Rad, USA) and UV-crosslinking at 1200 kDj/cm^2^ using UV Stratalinker 1800 (Stratagene, USA). After cross-linking membranes were pre-hybridised in ExpressHyb Hybridization Solution (Clontech, USA) at 40 °C for 1 h. The radioactive probes were prepared by end labelling 10 pmoles of DNA oligonucleotide complementary to the sfRNA (Supplementary table 7) with [γ-^32^P]-ATP (Perkin-Elmer, USA) using T4 polynucleotide kinase (NEB, USA). Unincorporated nucleotides were then removed by gel filtration on Illustra MicroSpin G-25 Columns (GE Healthcare, USA) and hybridisation was performed overnight at 40 °C in ExpressHyb Hybridisation Solution (Clontech, USA). Membranes were then rinsed and washed at 40 °C 4 x 15 min with a wash buffer containing 1% sodium dodecyl sulphate [SDS] and 1x saline-sodium citrate (SSC). Washed membranes were exposed to a phosphor screen (GE Healthcare, USA) overnight. Phosphor screens were scanned using Typhoon FLA 7000 Imager (GE Healthcare, USA).

### Viral cytotoxicity assay

Vero cells were seeded into 96-well plates at a density of 10^4^ cells per well. At 24 h after seeding, serial 0.5-log10 dilutions of virus samples were prepared, and 100 ul of each dilution were used to infect the cells. Eight wells on each plate were left uninfected (mock), to be used as control. Infection was performed by incubation with the inocula for 24 h, which were then replaced with 100 ul of DMEM medium supplemented with 2% FCS. At 3 dpi, cytopathic effects were determined based on cellular adenosine triphosphate levels using Viral ToxGlo Assay (Promega, USA) according to the manufacturer’s instructions. Luminescence was measured on a DTX880 Multimode Detector (Beckman Coulter). Percentage survival was determined as the percentage of luciferase activity (luminescence value) in infected cells compared to uninfected control, and cytopathic effect (CPE) was calculated as 100% – (% survival). The experiment was performed in duplicate. Data was fitted into sigmodal curves using GraphPad Prism v8.0.

### Virus growth kinetics

Cells were seeded at 10^6^ cells per well in 6-well plates and inoculated with wild type or mutated ZIKV at indicated MOIs by incubating for 1h with 200 μl of virus inoculum. Incubations were performed at 37° C, then inoculum was removed, cells were washed three times with PBS and overlayed with 2 ml of their relevant culture medium supplemented with 2% FBS. At time point zero, 100 µl of culture medium was immediately collected from the wells, and infected cells were then incubated for 3 days. Culture fluid samples (100 µl) were then harvested at 24, 48 and 72 hours post infection and subjected to focus-forming assay to determine the virus titres, from which growth curves were plotted.

### Animal Experiments

AG129 mice, which lack receptors for interferon α/β and γ, were obtained from animal facility at the Australian Institute for Bioengineering and Nanotechnology of the University of Queensland. The 5-6 weeks old AG129 mice (mix gender) were injected via intraperitoneal (i.p.) route with 10^3^ FFU/mouse of WT or sfRNA-deficient ZIKV_MR766_. Signs of encephalitis were monitored and scored as previously described^55^. Mice were monitored three time a day for signs of neurological disease and all mice with clinical score of three or more were immediately euthanized within 30 minutes by CO_2_ suffocation and cervical dislocation. Blood specimens were collected at indicated days post infection from the caudal vein, incubated at room temperature for 30 min and centrifuged at 10,000 x g at 4°C for 20 min to separate the serum.

For the pregnancy experiment, ten-week-old IFNAR1^−/−^ C57BL/6 pregnant mice were infected via subcutaneous injection with WT, xrRNA1’ or xrRNA2’ mutant of ZIKV_Natal_ at the dose of 10^4^ or 10^6^ FFU/mouse via subcutaneous injection. Infection of pregnant IFNAR^−/−^ dams was undertaken at E12.5. Mice were then monitored for 5 days and blood samples were collected daily as described above. At 5 days after infection (E17.5) dams were euthanized, foetuses weighted and photographed. Fetal heads and placenta were then processed for determination of tissue titers. One half of each tissue sample was homogenised in 1ml DMEM medium containing 2% FBS supplemented with Penicillin/Streptomycin (Gibco, USA) for 5min at 30Hz using a Tissue Lyser II (Qiagen, USA). Homogenates were then centrifuged at 10000 x g for 5 min at 4 °C and supernatants were collected for virus titration. Viral loads in mouse serum and tissue homogenates were determined by focus-forming assay on C6/36 cells using a human recombinant monoclonal antibody against ZIKV E-protein (hZ67).

### Human embryonic stem cells culture and generation of iPSC-derived human brain organoids

Human embryonic stem cells GENEA022^56^ (provided by Genea Biocells) were cultured on ECM Gel from Engelbreth-Holm-Swarm murine sarcoma (Sigma Aldrich Pty Ltd, Cat. # E1270-5X10ML) in mTeSR medium (Stem Cell Technologies, Cat. #85851). Cortical organoids were generated by an optimized protocol^57^ where patterned embryoid bodies were expanded for four days in N2 medium: DMEM/F12 (Gibco, Cat. #11320-33), 1% N-2 supplement (Gibco, Cat. #17502-048), 2% B-27 supplement (Gibco, Cat. # 17504044), 1% MEM Non-Essential Amino Acids (Gibco, Cat. #11140-050), 1% penicillin/streptomycin (Gibco, Cat. #15140148), 0.1% β-mercaptoethanol (Gibco, Cat. #21985-023) with daily supplementation of bFGF, 20 ng/mL (R&D, Cat. #233-FB-01M). Embryoid bodies were then embedded in 15 ul of Matrigel (Stem Cell Technologies, Cat. #354277) and media changed to 1:1 mixture of Neurobasal medium (Invitrogen, cat. #21103049) and DMEM/F12 medium supplemented with 1:200 MEM-NEAA, 1:100 Glutamax (Invitrogen, cat. #35050-038), 1:100 N2 supplement, 1:50 B27 supplement, 1% penicillin-streptomycin, 50 μM 2- mercaptoethanol and 0.25% insulin solution (Sigma, cat. #I9278). Fresh media was replaced thrice a week.

### Infection of iPSC-derived human brain organoids

Cerebral organoids at day 15 were utilized for viral infection. A virus inoculum titre of 10^4^FFU per 50 μl of each virus was added to a single organoid-containing well of a round-bottom, ultra-low-attachment 96-well plate and incubated at 37 °C O/N. Each ZIKV-infected organoid was then transferred to a single well of a 24-well plate containing 500 μl of ND medium. To determine viral growth kinetics in infected organoids, 160 μl of culture supernatant was harvested from each well at the indicated timepoints and then replaced by the same amount of fresh culture medium. Harvested culture fluids were titred by focus-forming assay. Three biologically independent organoids per virus were used. Infected organoids were imaged by dark-field microscopy with ×4 magnification using a Nikon Eclipse TE200 inverted microscope.

### Immuno-histological analysis of iPSC-derived human brain organoids

Organoids were fixed in 4% PFA for 1h at RT and washed three times with 1X phosphate buffer saline (PBS) at RT. Fixed organoids were then immersed in 30% sucrose in PBS at 4 °C. Organoids were allowed to sink before embedding in a solution containing 3:2 ratio of Optimal Cutting Temperature (O.C.T) and 30% sucrose. Embedded organoids were sectioned at 12-µm thickness with a Thermo Scientific™ CryoStar™ NX70 Cryostat. Organoid sections were air dried and washed three times for 10 minutes at RT followed by blocking and permeabilizing for 60 mins with a solution containing 3% bovine serum albumin (Sigma, Cat. A9418-50G) and 0.1% Triton X-100 in 1XPBS. Afterwards, sections were incubated with primary antibodies overnight at 4 °C and washed three times with 1X PBS for 10 minutes each at RT. Primary antibodies were anti-ZIKV E-protein 4G2 antibody used in 1:100 dilution; anti-NeuN (Millipore, ABN78, 1:500); anti-cleaved caspase 3 (Cell Signalling, 9661, 1:500); anti-Sox2 (Cell Signalling, D9B8N, 1:500). Secondary fluorophore-conjugated antibodies were then added for 1 hour at RT. All sections were incubated with Hoechst 33342 (Invitrogen, Cat. #H3570) for nuclei detection. Images were acquired using a Zeiss Axio Scan Z1 based in the School of Biomedical Sciences Imaging Facilities at the University of Queensland. The number of positive cells per organoid for the indicated markers was analysed by the imaging software CellProfiler, using the same pipeline for each sample in the same experiment.

### Caspase 3/7 activity assay

ReNcell human neural progenitor cells were seeded at the density of 5×10^5^ cells per well in 6-well plates coated with Matrigel Matrix Basement Membrane (Corning, USA) diluted 1:100 in PBS. Next day cells were infected at MOI=1. At 72 hpi culture media was removed, cells were washed twice with PBS and dislodged by incubation with 200 ul/well of Accutase Cell Detachment Solution (Innovative Cell Technologies, USA). Incubation was performed for 5 min at 37 °C, then digestion was terminated by addition of 200ul 0.1% trypsin inhibitor solution (Sigma-Aldrich, USA), cells were resuspended in 1ml PBS and then pelleted by centrifugation at 400 x g for 5 min. Cells were resuspended in 500 ul PBS and 50 ul of suspension was transferred into opaque 96-well plate for caspase assay. Caspase 3/7 activity was then assessed using Caspase-Glo 3/7 Assay System (Promega, USA) according to the manufacturer’s recommendations. Luminescence was measured using CLARIOstar Plus microplate reader (BMG Labtech, Germany).

### Next Generation Sequencing and bioinformatics analysis

RNA samples from three biological replicates of BeWo cells infected with WT or xrRNA2’ mutant ZIKV_MR766_ or mock-infected were subjected to poly(A) RNA enrichment and library preparation using TruSeq 2 Library Preparation Kit (Illumina, USA). Libraries were sequenced on an Illumina NextSeq 500 instrument using NextSeq 500/550 High Output Kit v2.5, 1×75 cycles (Illumina, USA). Image analysis was performed in real time by the NextSeq Control Software and Real Time Analysis running on the instrument computer. Quality control of raw sequencing data was performed using FastQC software v.0.72. Data was then trimmed to remove PCR primers, adapters and short reads using TRIMMOMATIC v.0.36.6 with the following settings: ILLUMINACLIP: TruSeq2-SE:2:30:10 LEADING:32 TRAILING:32 SLIDINGWINDOW:4:20 MINLEN:25 and subjected to another quality analysis with FastQC. Trimmed reads were mapped to the human genome assembly hg38 using HISAT2 v.2.1.0 allowing 1 mismatch. Feature counting was performed using featureCounts v1.6.4 with counting mode set to “Union”, strand to “Unstranded”, feature type was “exone” and ID attribute was Gene_ID. Genome FASTA and GFF3 files were obtained from NCBI Gene Bank. RNA-Seq data generated in this study is available in Gene Expression Omnibus database with accession number.

Differential gene expression analysis was performed using edgeR v.3.24.0. Low abundance reads (<1 cpm) were removed from the data set and data was normalised to library sizes and composition bias using trimmed mean of M-values (TMM) method. Normalised data was subjected to the multi-dimensional scaling analysis and used to build quasi-likelihood negative binomial generalized log-linear model. The quasi-likelihood F-test (glmQLFTest) was then applied to the contrasts WT-Mock, xrRNA2’-Mock and (WT-Mock)-(xrRNA2’-Mock). Genes were considered differentially expressed if FDR-corrected P-values were <0.05. Gene expression data were plotted using ggplot2 v.3.3.2.

Gene ontology and KEGG pathway enrichment analyses were performed using Database for Annotation, Visualization and Integrated Discovery (DAVID) v6.8. Enrichment data was then combined with expression values and z-scores were calculated using the R package GOplot v.1.0.2 and plotted using ggplot2 v.3.3.2. Heatmaps were generated using heatmap.2 function of R-package gplots v3.1.1. Gene interactions networks were reconstructed using Cytoscape v3.8.0. Genetic interactions were identified using GeneMANIA Cytosacape Plug-in. Betweenness centrality values represent the numbers of shortest paths that pass through each nod in the network and were calculated using “Analyze Network” function of Cytosacpe.

### Quantitative RT-PCR (qRT-PCR)

First strand cDNA synthesis was performed using qScript cDNA SuperMix (Quantabio, USA) according to manufacturer’s instructions. Quantitative PCR was then conducted using QuantStudio 6 Flex Real Time PCR Instrument (Applied Biosystems, USA). The cDNA was diluted 1:10 and 5 µl of the solution were used as template for qRT-PCR with SYBR Green PCR Master Mix (Applied Biosystems, USA). PCR was performed in 20 µl of a reaction mix containing 10 pmoles of forward and reverse PCR primers for each transcript (Supplementary table 7). Thermal cycling conditions were the following: 95 °C for 5 min and 40 cycles of 95 °C for 5 s and 60 °C for 20 s, followed by melting-curve analysis. Gene expression levels were determined by *ΔΔ*Ct method relative to mock and normalised to *GAPDH*. Viral genomic RNA levels were determined using standard curve approach by comparing Ct values of the samples to Ct values observed in amplification of serial dilutions (10^2^ – 10^8^ copies/reaction) of a PCR-amplified and purified ZIKV genomic fragment. For each experiment, RNA from 3 biological replicates was used and PCR amplification of each cDNA sample was performed in triplicate. Negative controls were included for each set of primers.

### Interferon treatment

To examine STAT1 phosphorylation in infected cells Vero or HTR-8 cells were seeded in 24-well plates at a density of 2×10^5^ cells per well. Next day cells were infected with WT or xrRNA2’ mutant of ZIKV_MR766_ at MOI=5. At 48 hpt, cell culture medium was replaced with 1ml fresh medium containing 10^4^ units of human IFN-*α*2 (Abcam, UK) or 100ng/ml of human IFN-*λ*1 (PeproTech, USA). Vero cells were incubated with IFN-*α*2 for 20min and HTR-8 cells were incubated with IFN-*λ*1 for 1h. Immediately after incubations cells were lysed for western blotting or fixed for immunofluorescent protein detection.

### Western blotting

Cells were lysed in 100 μl per well of 24-well plate of Bolt LDS Sample Buffer (ThermoFisher Scientific, USA) supplemented with Bolt Sample Reducing Agent (ThermoFisher Scientific, USA) and Protease Inhibitor Cocktail (Sigma-Aldrich, USA). Lysates were sonicated using Branson Digital Sonifier 450 (Marshall Scientific) at 10% output for 15 sec with pulse on for 3 sec and off for 1 sec. Lysates were then incubated at 95°C for 5 minutes, cooled on ice and loaded into the Bolt 4-12% Bis-TRIS gels (ThermoFisher Scientific, USA). Electrophoresis was performed for 1 hour at 170V in Bolt MES SDS Running Buffer (ThermoFisher Scientific, USA). Proteins were then electroblotted onto a nitrocellulose membrane using iBlot2 Dry Blotting System (ThermoFisher Scientific, USA) and iBlot 2 Transfer Stack (ThermoFisher Scientific, USA). Membranes were blocked in Clear Milk Blocking Buffer (Pierce, USA) for 1 hour, incubated with primary antibodies overnight at 4°C, then washed 4 x 5 min with TBS-T, incubated with HRP-conjugated secondary antibodies for 1 hour at RT and washed 4 x 5 min with TBS-T. Detection of HRP activity was performed using SuperSignal West Pico PLUS Chemiluminescence Substrate (Pierce, USA) and signal was visualised using Amersham Imager 600. Blot densitometry was conducted using ImageJ Fiji v.2.1.0. The primary and secondary antibodies and their dilutions are listed in Supplementary table 8.

### Immunofluorescence assay

Cells for IFA were seeded on glass cover slips contained in 24-well plate. After IFN-treatment cells were washed once with PBS and fixed with 500ul per well of ice-cold 100% methanol at -20 °C for 20 min. Fixed cells were rinsed twice with 500 ul TBS and incubated with 500 ul of 50mM Glycine for 30 min at RT to quench autofluorescence. Cells were then washed 3 x 5 min with 500 ul TBS, blocked for 1h at RT with 500 ul of 1% BSA in TBS and incubated at 37 °C for 1h with 250ul of primary antibodies diluted in blocking solution. The mixture of antibodies contained rabbit monoclonal antibody against phospho_Tyr701_-STAT1 (#9167, Cell Signalling Technologies, USA) diluted 1:200 and mouse monoclonal antibody 4G2 against flavivirus E-protein diluted 1:50. Cells were then washed 3 x 5 min with 500 mml per well of TBS and incubated at 37°C for 1h with 250ul of a mixture containing goat anti-rabbit IgG (H+L) highly cross-adsorbed secondary antibody conjugated with Alexa Fluor plus 488 (Invitrogen, USA) and goat anti-mouse IgG (H+L) highly cross-adsorbed secondary antibody conjugated with Alexa Fluor 594 (Invitrogen, USA), each diluted 1:500 in blocking solution. After the incubation cells were washed 3 x 5 min with 500 ul per well of TBS and cover slips were mounted using ProLong Gold antifade reagent with DAPI (Invitrogen, USA). Imaging was performed using Zeiss LSM710 confocal microscope.

### In vitro transcription and transfection of RNA

For *in vitro* transcription plasmids containing sequence of interest under control of T7 promoter were linearized by restriction digest and 1 ug of purified digested DNA was used as a template for the reaction performed using MEGAscript T7 Transcription Kit (Ambion, USA) as specified by the manufacturer and purified by LiCI precipitation. For generation of ZIKV sfRNA and GFP RNA fragment, fluorescein-labelled RNA was prepared in parallel using Fluorescein RNA Labelling Mix (Roche, Switzerland) and mixed with unlabelled RNA in 1:10 ratio to serve as a tracer. RNA was then incubated with 10U RppH (NEB, USA) and 1U XRN-1 (NEB, USA) in NEB3 Buffer for 1h at 37°C. RNA was then purified by Phenol:Chlorophorm extraction. All in vitro transcribed RNAs were analysed by electrophoresis in 1.5% formaldehyde agarose gel and RNA concentration was determined on Nanodrop ND-1000 spectrophotometer (ThermoFisher Scientific, USA). RNA was transfected into cells using in-suspension protocol^53^.

### RNA affinity pull-down

To generate the plasmids for in vitro transcription of sfRNA and negative control RNA fused with four streptavidin affinity tags (4xS1m), the aptamer sequences followed by ZIKV sfRNA sequence or GFP gene fragment were inserted under T7 promoter into pUC19 vector. The DNA fragments containing T7 promoter with 2xS1m and 2xS1m alone (T7_2xS1m_SacI_BamHI_F/R and 2xS1m_BamHI_XbaI_F/R in Supplementary table 7), were obtained as two sets of complementary single-stranded Ultramer® DNA oligos (IDT, USA). These oligos were designed to form terminal overhangs compatible with restriction enzyme cleavage sites within pUC19 vector upon annealing. Prior to cloning, 2 nmols of each of the complementary oligonucleotides were annealed by incubation at 95°C for 5 minutes, followed by gradual cooling to room temperature in 50 ul of Duplex Buffer (IDT, USA). Annealed oligos were gel-purified from 2% agarose gel and T7_2xS1m_SacI_BamHI DNA fragment was ligated with SacI+BamHI-digested pUC19 plasmid using T4 DNA ligase (NEB). The resulted plasmid was digested with BamHI and XbaI and ligated with 2xS1m_BamHI_XbaI dsDNA oligo to generate pUC19-T7-4xS1m construct. The ZIKV sfRNA sequence and GFP gene fragment were PCR amplified from pUC19-ZIKV-F4 ^14^ and pMYC-GFP (Addgene, #42142) plasmids respectively using primers HindIII_ZIKV_sfRNA and XbaI_GFP (Supplementary table 7) and PrimeSTAR GXL Polymerase (Takara, Japan). Cycling conditions were as follows: 35 cycles of 98 °C - 10 sec, 55°C - 30 sec, 68°C – 3 min 30 sec, followed by a final extension at 68°C for 5 min. PCR products were gel-purified, digested with XbaI and HindIII (NEB, USA) and ligated into pUC19_4xS1m plasmid digested with the same enzymes. The resulted constructs pUC19_sfRNA_4xS1m and pUC19_GFP_4xS1m as well as all intermediate plasmids were Sanger sequenced and shown to conform the desired design and sequence.

RNA affinity purification was performed using optimized S1m streptavidin RNA-aptamer^58^ and previously described protocol with minor modifications^30^. A549 cells were seeded in 10 x T175 flasks at a density of 1×10^7^ cells per flask and infected with ZIKV_MR766_- xrRNA2’ at an MOI = 0.5. At 48 hours post-infection (hpi), the cells were washed once with ice-cold DEPC-treated PBS, harvested by scraping in 10 ml DEPC-PBS per flask, pelleted by centrifugation at 300 x g for 5 min and washed twice in DEPC-PBS. Cells were then lysed by incubation on ice for 30 min in 2ml of SA-RNA lysis buffer (20 mM Tris-HCl pH 7.5, 150 mM NaCl, 1.5 mM MgCl2, 2 mM DTT, 2mM vanadylribonucleosid complex RNase inhibitor (NEB, USA)), supplemented with 1x Protease Inhibitor Cocktail (Sigma-Aldrich), 1x PhosphoSTOP Phosphatase Inhibitor Cocktail (Sigma-Aldrich, USA) and 1% NP-40 (Sigma-Aldrich, USA). The cell lysates were then sonicated using Branson Digital Sonifier 450 (Marshall Scientific) at 10% output for 15 sec with pulse on for 3 sec and off for 1 sec. Lysates were cleared by centrifugation at 16000 x g for 15 min at 4 °C and split in halves for incubation with sfRNA bait and GFP bait.

The in vitro transcribed, aptamer-containing RNA was mixed with SA-RNA lysis buffer to make up a final volume of 50 ul. RNA was renatured by incubation at 56°C for 5 minutes, 37°C for 10 min and RT for 2 min, then chilled on ice. For RNA to bead coupling, 100 ul of Streptavidin Sepharose High Performance bead-slurry (GE Healthcare, USA) was first equilibrated by washing three times with 500 ul of SA-RNP lysis buffer and collected by centrifugation at 100 x g for 2 min, then resuspended in 100 ul of SA-RNP lysis buffer supplemented with 80U RNAsin (Promega, USA) and incubated with 30 ug of renatured RNA with overhead rotation at 4°C for 2.5h. The A549 cell lysates were pre-cleared by incubation with 100 ul of equilibrated SA-beads for 2.5h at 4°C with overhead rotation. Beads were then removed by centrifugation at 16000 x g for 1 min, lysates were supplemented with 40U RNasin (Promega, USA) and incubated with sfRNA_4xS1m or GFP_4xS1m- coupled SA-beads at 4°C for 3.5h with overhead rotation. Then, the protein-bound SA-beads were collected by centrifugation at 100 x g for 2 min, and washed five times with SA-wash buffer (20 mM Tris-HCl pH 7.5, 300 mM NaCl, 5 mM MgCl2, 2 mM DTT, 1xVRC and 1x Protease Inhibitor Cocktail (Sigma-Aldrich)), followed by a final wash with SA-wash buffer supplemented without supplements. Bound proteins were eluted from the beads by incubation with 100ul of 8M Urea / 50mM Ammonium Bicarbonate (ABS) for 15min with overhead rotation. Beads were removed by centrifugation at 3800 x g for 1 min and the elutes were collected.

### Mass spectrometry

Pull-down elutes were centrifuged at 16,000 x g for 10 min to remove insoluble materials and protein concentrations were determined using Pierce BCA Protein Assay Kit (Pierce, USA). Samples containing 10 ug of protein were transferred to an Amicon Ultra 10k 0.5 ml Centrifugal Filter column (Amicon, USA) and centrifuged at 14,000 x g for 40 min. The column was washed with 500 ul of wash solution (8M urea, 50 mM ammonium bicarbonate (ABC)) and centrifuged at 14,000 x g for 4 min. The on-column reduction of the proteins was then performed in 200 ul wash solution containing 5mM DTT at 56 °C for 30 min. Then iodoacetamide (IAA) was added to the solution to a final concentration of 25 mM and alkylation was performed at RT in the dark for 30 min. reaction was terminated by adding DTT to a final concentration of 10 mM and reaction solution was removed by centrifugation at 14,000 x g. Proteins were then resuspended in 100 ul of 50 mM ABC and digested overnight at 37 °C by using of 0.2 ug trypsin per 10 ug of protein. To collect the peptides, the column was transferred to another collection tube and centrifuged at 14,000 x g, RT for 40 min. 50 ul of 0.5 M NaCl was then added to the column and centrifuged through to ensure the complete elution of the peptides. The elutes were combined and the peptides were further purified using C18 ZipTip Pipette Tips (Millipore, USA) as per manufacturer’s instructions.

Peptides were separated using reversed-phase chromatography on a Shimadzu Prominence NanoLC system. Using a flow rate of 30 µl/min, samples were desalted on an Agilent C18 trap (0.3 x 5 mm, 5 µm) for 3 min, followed by separation on a Vydac Everest C18 (300 A, 5 µm, 150 mm x 150 µm) column at a flow rate of 1 µl/min. A gradient of 10-60% buffer B over 45 min where buffer A = 1% ACN, 0.1% FA and buffer B = 80% ACN, 0.1% FA was used to separate peptides. Eluted peptides were directly analysed on a TripleTof 5600 instrument (ABSciex) using a Nanospray III interface. Gas and voltage settings were adjusted as required. MS TOF scan across m/z 350-1800 was performed for 0.5 sec followed by information-dependent acquisition of the top 20 peptides across m/z 40-1800 (0.05 sec per spectra).

Reference protein sequences for human and ZIKV_MR766_ proteins were obtained from SwissProt database. MS data were converted to mgf format and searched using Paragon method and ProteinPilot software (Sciex, USA). Search settings were the following: trypsin was set as the enzyme, the instrument was set to TripleTof 5600, cys modifications were set to iodoacetamide. Peak areas were quantified using in PeakVeiw (Sciex, USA) using the SWATH Acquisition MicroApp (Sciex, USA). Protein enrichment analysis was performed on peak area data using Limma v3.46.0 and R v3.6.3.

### Generation of expression plasmids

To generate the expression plasmids that produce ZIKV NS5 with sfRNA, and GFP with ZIKV sfRNA, DNA fragments containing Kozak sequence followed by the corresponding ORF, ZIKV 3’UTR and HDV ribozyme were cloned into pcDNA3.1(+) expression vector. The fragments were obtained as synthetic custom genes cloned in pUCIDT vector (IDT, USA). To introduce the restriction sites compatible with pcDNA3.1(+), fragments were PCR amplified from the plasmids using PrimeSTAR GXL Polymerase (Takara, Japan) and Nhel_IDT_F and EcoRI_ITD_R primers (Supplementary table 7), which are complementary to the pUCIDT vector backbone immediately upstream and downstream of the insert. The cycling conditions were as follows: 98°C for 2 minutes, 35 cycles of 98°C for 10 seconds, 55°C for 30 seconds, and 68°C for 3 minutes and 30 seconds, followed by a 5-minute extension at 68°C. The two PCR amplicons were gel-purified, digested with NheI and EcoRI, and ligated with pcDNA3.1(+) vector that was cut with the same enzymes.

To generate the plasmids pcDNA3.1(+)-ZIKV_NS5_3’UTR-Mut and pcDNA3.1(+)- GFP_ZIKV_3’UTR that respectively produce ZIKV NS5 without sfRNA, and neither NS5 nor sfRNA (negative control), PCR directed site-specific mutagenesis was used to introduce point mutations into two xrRNA regions of 3’UTRs within pcDNA3.1(+)-ZIKV_NS5_3’UTR and pcDNA3.1(+)-GFP_ZIKV_3’UTR plasmids. Mutagenesis was performed in two steps using Q5 site-directed mutagenesis kit (NEB, USA) as per manufacturer’s instructions. Briefly, the first round of PCR mutagenesis was conducted using xrRNA1_C22G primers (Supplementary table 7). The cycling conditions were as follows: 98°C for 30 seconds, 35 cycles of 98°C for 10 seconds, 55°C for 30 seconds, and 72°C for 4 minutes and 30 seconds, followed by a 2-minute extension at 72°C. The PCR-amplified plasmids were circularised, propagated in *E.coli* and used as templates for the second round of PCR mutagenesis, under the same cycling conditions with xrRNA2_C22G primers. All expression plasmids were deep-sequenced using Oxford Nanopore platform.

### Nanopore Sequencing

Nanopore sequencing of amplicons was performed as described previously^59^. The regions of the expression plasmids containing the entire inserts were PCR amplified using PrimeSTAR GXL Polymerase (Takara, Japan) and Nanopore_pCMV_F and Nanopore_BGH_R primers (Supplementary table 7). The amplicons were gel-purified and subjected to the preparation of a barcoded libraries using PCR Barcoding Expansion Kit (EXP-PBC001, Oxford Nanopore Technologies, UK). Purified barcoded DNA was subjected to End Repair/dA-tailing using NEBNext® Ultra™ II Module (NEB, USA) and ligated to the sequencing adapters using Ligation Sequencing Kit (SQK-LSK109, Oxford Nanopore Technologies, UK). The prepared library was quantified using the Qubit™ dsDNA HS Assay Kit (ThermoFisher Scientific, USA), and 20 fmol of the final library was loaded to a MinION Flongle flow cell (Oxford Nanopore Technologies, UK). Sequencing runs were conducted using the MinKNOW software (Oxford Nanopore Technologies, UK). Base-calling and adapter trimming of the fast5 files containing sequencing reads were conducted using guppy_basecaller v 3.3.0+ef22818 (Oxford Nanopore Technologies, UK). The fastq files containing the reads were used for reads mapping against the reference plasmid sequence using the Bowtie2 tool (Galaxy Australia). The binary alignment file was visualized using Integrative Genomics Viewer 2.8 (Broad Institute, USA).

### Plasmid transfection

HEK293T cells were seeded in a 24-well plate at a density of 1.5 x 10^5^ cells per well, and transfected with 0.14 pmol of plasmid DNA using 2.5 μl per well of Lipofectamine 2000 (Invitrogen, USA) according to the manufacturer’s instructions. At 48 hpt, cells in each well were treated with 50 units of human IFN-*α*2 (Abcam, UK) for 15 min or left untreated as a control, then washed with PBS and lysed for western blotting.

### Statistical analysis

Statistical analysis was performed using Graph Pad Prism software version 8 and R Studio version 0.99.893. Details of statistical methods are described in figure legends and supplementary tables.

### Ethics statement

All mouse work was conducted in accordance with the “Australian code for the care and use of animals for scientific purposes” as defined by the National Health and Medical Research Council of Australia. Mouse work was approved by the QIMR Berghofer Medical Research Institute animal ethics committee (P2195) and University of Queensland animal ethics committee (AE31401).

All experiments with human stem cells were carried out in accordance with the ethical guidelines of the University of Queensland and with the approval by the University of Queensland Human Research Ethics Committee (Approval number-2019000159). A commercially available hESC line was used. The manufacturer reported that karyotyped embryo cells were fully consented for the development of stem cells by all responsible people through an informed consent process (signed de-identified consent form can be provided upon request). Donors have received no payment of other benefits for their donation. Donated embryos were originally created by assisted reproduction technology (ART) for the purpose of procreation. Embryos were identified as unsuitable for implantation, biopsy or freezing due to abnormal development. Embryonic outgrowths were developed for consented clinical investigation studies.

### Data availability

All data are available within the article, supplementary files and upon request from the authors. RNA sequencing data sets are deposited in NCBI Gene Expression Omnibus database with accession number GSE171648.

## Acknowledgments

We acknowledge the excellent support of Amanda Nouwens and Peter Josh from School of Chemistry and Molecular Biosciences (The University of Queensland) in mass spectrometry and proteomics data analysis. We are thankful to Maria Kasherman and Shaun Walters from the School of Biomedical Sciences Imaging and Analytical facilities (The University of Queensland) for technical support. We are thankful to Prof. Roy Hall (The University of Queensland), Prof. Andres Merits and Dr Eva Žusinaite (University of Tartu), Prof. Bryan Day and Dr. Ulrich Baumgartner (QIMR Berghoffer Medical Research Institute) for providing antibodies and cell lines. Next generation sequencing was performed at ACE Sequencing (The University of Queensland). Sanger sequencing was performed by Australian Genomics Research Facility. Work was funded by NHMRC grants APP1127916 and APP1059794 to AAK. JA was supported by a University of Queensland Early Career Researcher Grant (UQECR2058457), an NHMRC Ideas Grant (APP2001408), a Jérôme Lejeune Postdoctoral Fellowship and a Brisbane Children’s Hospital Foundation grant (Project-50308). EW was supported by the NHMRC (grants APP1138795, APP1127976, APP1144806 and APP1130168), BrAshA-T Foundation, Perry Cross Spinal Research Foundation and an ARC Discovery Project (DP210103401).

## Author contributions

A.Sl., X.W., H.C., J.A., M.F., K.Y., F.J.T., A.A.A., R.B., Y.X.S., J.D.J.S., N.P. – experiments; A.Sl., A.Su. – bioinformatics; A.Sl., A.A.K. – conceptualization, experiment design; A.Sl., X.W., J.A., A.Sl. – data analysis; D.W. – critical reagents; E.W., A.Su., A.A.K. – project supervision; A.Sl., A.Su., A.A.K. – manuscript preparation, A.A.K. – funding acquisition.

